# Breakdown in the synaptic vesicle cycle defines early and reversible cortical pathogenesis in ALS

**DOI:** 10.64898/2026.08.21.746168

**Authors:** Zsofia I. Laszlo, Anna Sanchez-Avila, Anna McFarlane, Dinja van der Hoorn, Rebecca San Gil, Tara L. Spires-Jones, Thomas H. Gillingwater, Adam K. Walker, Christopher M. Henstridge

## Abstract

Synaptic failure is considered an early driver of Amyotrophic Lateral Sclerosis (ALS), yet identifying the molecular events initiating synaptic decline remains challenging in end-stage human tissue. Here, we exploit the late involvement of the primary visual cortex (Brodmann Area 17 (BA17)) to investigate early disease-associated changes in human ALS. Structural analyses revealed neuropil compaction, presynaptic terminal shrinkage, and synaptic degeneration despite preservation of local neuronal populations. Deep synaptoneurosome proteomics identified a regional signature characterised by disruption of presynaptic vesicle cycling, which closely resembles early pathological changes observed in the inducible human TDP-43 rNLS8 mouse model. Importantly, suppression of TDP-43 expression *in vivo* restored these proteomic alterations, highlighting recovery of presynaptic vesicle machinery within preserved synaptic structures. Together, these findings reveal early synaptic pathology as a distinct and potentially reversible stage of ALS neurodegeneration.

## Main

Amyotrophic Lateral Sclerosis (ALS) is a fatal neurodegenerative disease characterised by progressive muscle wastage and paralysis. Motor neuron degeneration in both the brain and spinal cord underlies the physical symptoms, yet growing evidence suggests ALS is a multisystem disorder with widespread CNS dysfunction. Prior to neuronal loss, early synapse dysfunction and breakdown is observed as a convergent feature across ALS models and is evident in human CNS post-mortem tissue^1–3^.

Cortical hyperexcitability occurs early, even presymptomatically in genetic cases^4^, and contributes to neuronal vulnerability via synaptic excitotoxicity. Multiple synaptotoxic processes contribute, including altered glutamatergic signalling, reduced inhibitory transmission, fractured synaptic scaffolding, disrupted vesicle cycling and altered dendritic spine maintenance^3,5,6^, highlighting early failures of key synaptic processes. Furthermore, ALS patient-derived neurons display similar synaptotoxic phenotypes, which ultimately develop into synaptic breakdown, neuronal hypoexcitability and cell death, recapitulating key features underlying clinical presentation^5,7^.

The involvement of extra-motor regions in ALS is supported by the frequent occurrence of cognitive and/or behavioural impairments which overlap with frontotemporal dementia (FTD)^8^. Synaptic abnormalities have been identified in frontal and limbic regions of these patients, while ALS models carrying mutations in *SOD1, TARDBP, C9orf72 and FUS* demonstrate early synaptic loss, altered excitatory–inhibitory balance and circuit dysfunction beyond motor regions^9–11^. These findings suggest that synaptic vulnerability is a fundamental feature of ALS pathogenesis rather than a consequence restricted to motor neuron degeneration.

Although sensory systems were historically considered unaffected in ALS, increasing evidence indicates involvement of the visual cortex (BA17). Neuroimaging studies have reported occipital cortical thinning, altered visual network connectivity and changes in visually evoked potentials in ALS patients^12^. More recently, SV2A PET imaging demonstrated reduced synaptic density within the occipital lobe, supporting the presence of synaptic degeneration beyond classical ALS-affected motor regions^13^. Consistent with these findings, pathological TDP-43 accumulation is frequently observed in non-motor cortical regions, including frontal, temporal and occipital areas, suggesting that ALS-associated proteinopathy extends across interconnected neural networks, in a temporal manner^14–16^. The stereotypical distribution of TDP-43 pathology has led to the hypothesis that misfolded TDP-43 may propagate through synaptically connected neuronal circuits in a prion-like manner; however, the mechanisms determining regional vulnerability and the earliest molecular consequences of this process remain unclear^17,18^. In particular, how early TDP-43 pathology contributes to synaptic dysfunction in vulnerable but non-degenerating regions remains poorly understood.

Temporally controlled models of TDP-43 proteinopathy provide an important experimental framework to investigate these early disease mechanisms. The doxycycline-controllable TDP-43 rNLS8 mouse model enables precise control of TDP-43 expression and the investigation of temporal molecular changes preceding overt neurodegeneration^19^. Following induction, cytoplasmic TDP-43 accumulation leads to a well-characterised progressive motor neuron degeneration, neuromuscular junction loss and motor impairment^19–21^. Importantly, this model captures early pathogenic events, including activation of stress responses, transcriptional dysregulation, synaptic impairment and neuroinflammation, before substantial neuronal loss occurs^22^.

To uncover the key disease-driven molecular changes at the synapse, we combined high-resolution human neuropathology and sophisticated synaptoneurosomal proteomics to deeply phenotype the human visual cortex. We reveal a shrinkage of presynaptic terminals, and a regional and disease-specific synaptic molecular signature, highlighting presynaptic vesicular breakdown as a key feature. To confirm this early synaptic phenotype, we used the inducible rNLS8 mouse, collected synaptosomes at several disease stages and performed the same proteomic analysis as for our human samples. Aligning the datasets revealed that our human data clustered with the early disease stage and highlighted presynaptic vesicular breakdown as central to early synaptic phenotypes in the model. The mice also underwent a recovery period where the transgene was switched off; importantly, we observed recovery of key synaptic proteins. Our findings suggest presynaptic vesicular breakdown is an early, yet reversible phenotype in ALS.

Collectively, we present a comprehensive and integrated synaptic protein resource across ALS models and human post-mortem material. These data highlight early mechanisms of synaptic breakdown and are fully accessible through our open-access web-based platform (https://henstridgelab.shinyapps.io/mnd-synaptic-proteomics/).

## Results

### Cellular and Subcellular Alterations in the ALS Visual Cortex

Neuropathological staging paradigms indicate that the primary visual cortex (BA17) represents one of the final anatomical milestones of cortical disease progression in ALS (**Fig. 1a**). This spatiotemporal lag provides a unique window for capturing early human disease mechanisms before the onset of widespread pathological protein deposition and tissue destruction.

**Figure 1.**
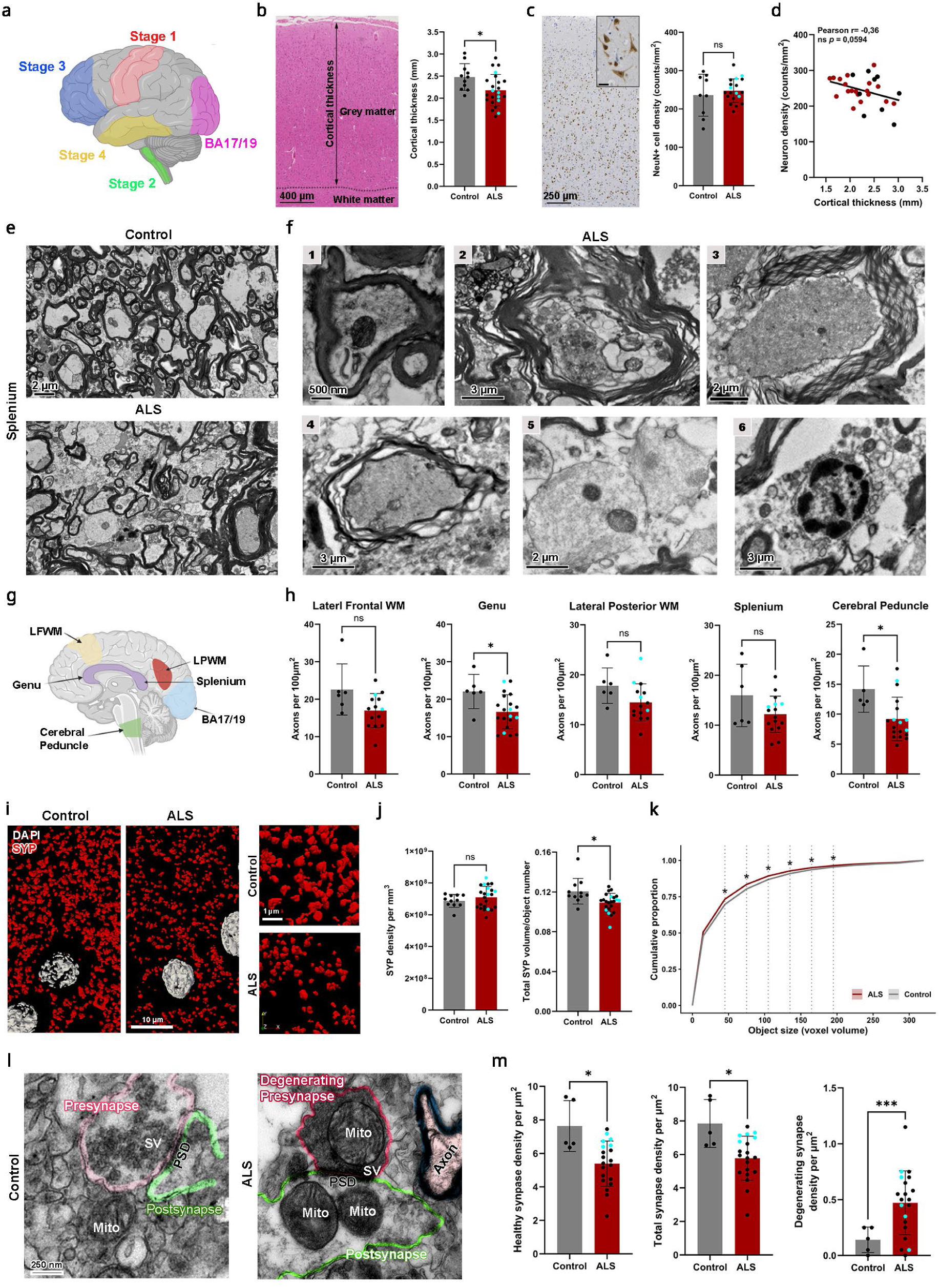
Cellular and subcellular alterations across the ALS primary visual cortex. **a**, Schematic of the amyotrophic lateral sclerosis (ALS) neuropathological staging paradigm, illustrating the sequential spread of phosphorylated TDP-43 (phTDP43) pathology from the primary motor cortex (Brodmann area 4; BA4) and brainstem motor nuclei (Stage 1), to the brainstem reticular formation precerebellar nuclei and thalamus (Stage 2), to the prefrontal/postcentral neocortex and striatum (Stage 3), and to the anteromedial temporal lobe and hippocampus (Stage 4). **b**, Representative image of H&E-stained cortex and quantification of grey matter atrophy in the ALS visual cortex. Ctrl n=11, ALS n=23, Welch’s t test p=0.0182. **c**, Representative images and quantification of neuron density based on NeuN staining. Ctrl n=9, ALS n=19, Welch’s t test p=0.575. **d**, Correlation analysis between cortical thickness and NeuN-positive neuron density, n = 28, Pearson r =-0.3606, p=0.0594. **e**, Electron micrographs showing disrupted white matter architecture and reduced myelin integrity in ALS compared to control tissue. **f**, Microstructural stages of myelinated axon breakdown (panels 1–6): (1) densely packed axon with enlarged mitochondria; (2) demyelinated axons with intracellular organelle accumulation; (3) advanced demyelination with loose myelin sheaths; (4) condensed axoplasm with thin myelin lamellae; (5) fully demyelinated axons; (6) condensed chromatin within glial cell nuclei. **g**, Schematic of examined brain regions: lateral frontal white matter (LFWM; BA6/8 level), anterior (genu) and posterior (splenium) corpus callosum, lateral posterior white matter (LPWM; BA39 level), and the cerebral peduncle (CP) of the corticospinal tract. **h**, Quantification of axonal coverage across the indicated white matter regions. Welch’s t test, LFWM: p=0.1066 Ctrl n=6, ALS n=14, Genu p=0.0335 Ctrl n=6, ALS n=20, Splenium p=0.2103 Ctrl n=6, ALS n=16, LPWM p=0.0826 Ctrl n=6, ALS n=15, Mann Whitney test, CP p=0.0066 Ctrl n=5, ALS n=18. Representative 3D-rendered array tomography images (**i**) and quantification (**j**) of synaptophysin (SYP) immunofluorescence density and object volume. SYP density: Welch’s t test p=0.2435, SYP volume: Welch’s t test p=0.0198, Ctrl n=11, ALS n=23. **k**, Cumulative distribution of SYP-positive object volumes in ALS and control visual cortex binned at 30-voxel intervals (two-sided Wilcoxon rank-sum test with Benjamini–Hochberg correction; * p < 0.05). Electron microscopy images (**l**) and quantification (**m**) of synapse loss and degeneration in ALS, defined by dense presynaptic axoplasm, enlarged mitochondria, and vacuolisation. Welch’s t test, total density p=0.0266, healthy synapses p=0.0253, degenerating synapses p=0.0008, Ctrl n=5, ALS n=20. Cyan symbols indicate *C9orf72*-positive cases; bar graphs show mean±SD.

We began by profiling phosphorylated TDP-43 (pTDP43) pathology; remarkably, 13 out of 23 ALS cases exhibited pTDP43-positive inclusions within BA17, although most were very rare (**Extended Data Fig. 1a,b**). In the majority, these visual cortex inclusions were found in donors that also had inclusions in the earlier-affected primary motor (BA4) and prefrontal (BA9) cortices, aligning with published pTDP43 staging paradigms^14,15^.

To evaluate macro-structural integrity, we quantified cortical grey matter thickness, revealing a modest but significant 15% thinning of the ALS BA17 cortex (**Fig. 1b**). This was not driven by significant neuron loss; instead, we observed a stable (slight 5% increase) density of NeuN+ve cells within the thinned ALS cortex (**Fig. 1c**). Intra-cohort correlation analysis revealed an inverse relationship between cortical thickness and neuronal density (**Fig. 1d**). Geometrically, this negative correlation provides evidence of a tissue-level compaction demonstrating that the macro-scale thinning of BA17 is likely driven by reduction of the surrounding neuropil prior to neuronal death.

Going upstream of this neuropil volume loss, we interrogated the underlying white matter tracts for evidence of axonal degeneration. Qualitative ultrastructural analysis of the splenium (the posterior corpus callosum tract projecting directly into BA17) revealed an array of degenerative axonal profiles and myelin fragmentation in ALS tissues (**Fig. 1e,f**). This was accompanied by striking cell loss (**Extended Data Fig. 1c,d**) and robust reactive astrogliosis (**Extended Data Fig. 1e**) within the underlying BA17 white matter. Intriguingly, white matter CD68+ve microglial reactivity was significantly reduced (**Extended Data Fig. 1f**), while the overlying grey matter lacked reactive astrogliosis or changes in microglial activation (**Extended Data Fig. 1g,h**). This regional compartmentalisation suggests that the grey matter in BA17 remains largely quiescent in terms of gliosis, despite underlying white matter pathogenesis.

High-resolution quantification revealed axonal depletion across several BA17-projecting long-range white matter tracts (**Fig. 1g,h**), establishing that upstream tract collapse may drive some of the downstream cortical thinning.

If incoming axonal tracts are dismantling, we reasoned that their synaptic terminals within the grey matter likely undergo a parallel reorganization. To resolve this at nanometer resolution, we deployed two complementary high-resolution imaging techniques. Quantitative array tomography (AT) revealed that the density of Synaptophysin-positive (SYP+ve) presynaptic terminals remained unaltered in ALS (**Fig. 1i,j, Extended Data Fig. 2**). However, within a significantly thinned cortical volume, a static terminal density mathematically suggests a loss of absolute synapse numbers across the cortical column.

AT volumetric analysis revealed that surviving SYP+ve puncta were significantly smaller in ALS donors, displaying a leftward shift toward a condensed size distribution (**Fig. 1j,k**). Direct ultrastructural analysis via transmission electron microscopy (TEM) confirmed a loss of healthy synapses, alongside a robust accumulation of electron-dense degenerating synaptic remnants (**Fig. 1l,m**).

Collectively, these data demonstrate that early pathogenesis in the human ALS cortex is defined by presynaptic terminal shrinkage and protein condensation that precedes neuronal cell death.

### Alterations in the ALS BA17 Synaptic Proteome

Having established physical evidence of a presynaptic phenotype, we next sought to identify the molecular cascades underlying this terminal decay. We isolated intact synaptoneurosomes from ALS and control BA17 donor tissues and processed individual fractions via label-free, data-independent acquisition (DIA) mass spectrometry (**Fig. 2a, Extended Data Fig. 3**). Because this biochemical approach isolates intact, re-sealed synaptic junctions observed alterations in the proteome reflect true cell-autonomous adjustments in terminal protein composition rather than mathematical artifacts driven by synapse loss.

**Figure 2.**
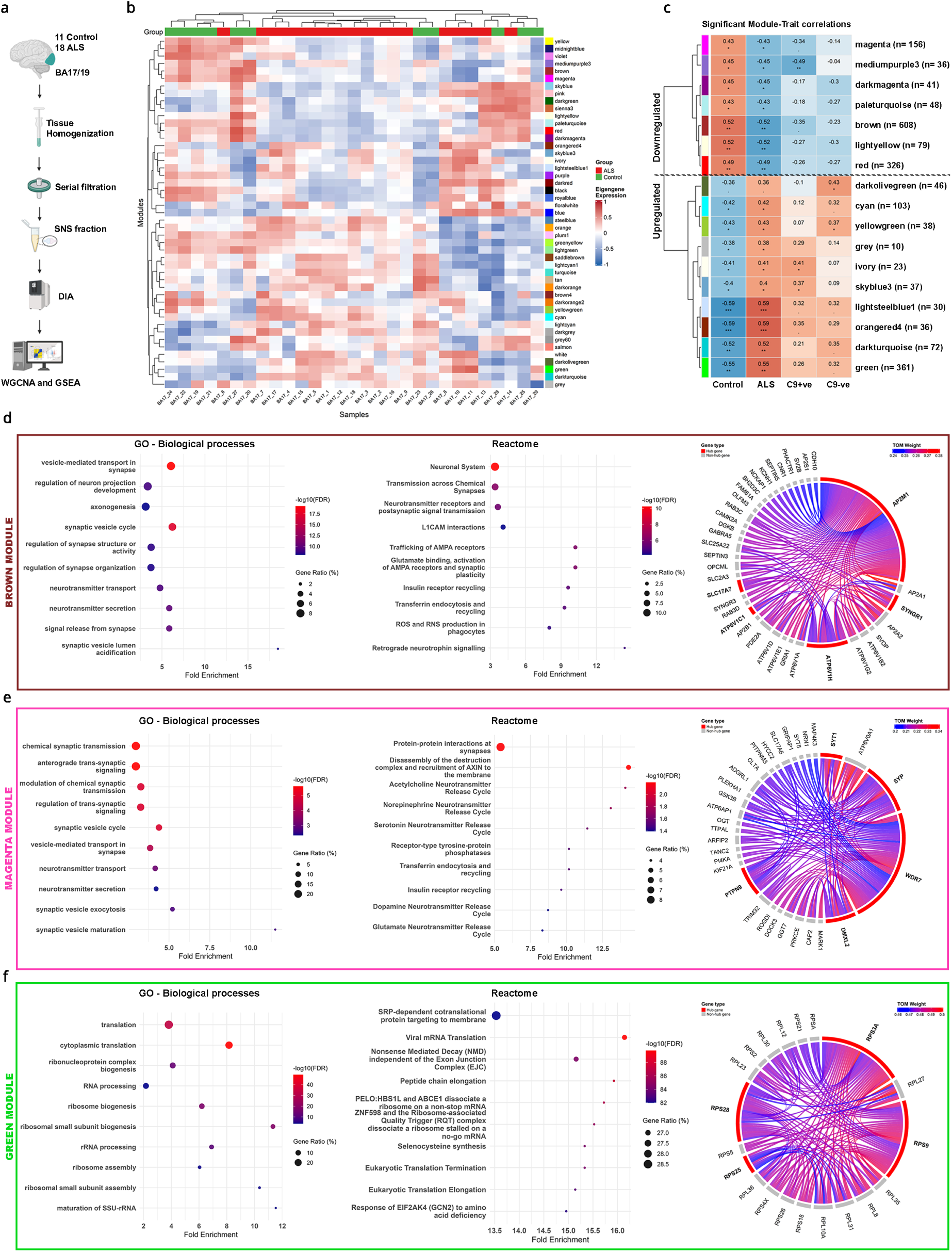
WGCNA identifies presynaptic dysfunction in the ALS visual cortex. **a**, Schematic representation of the analytical and computational workflow. **b**, Module eigengene heatmap displaying module expression profiles across individual samples and clinical traits. **c**, Hierarchical heatmap showing significant module–trait correlations calculated using biweight midcorrelation (bicor). Significance indicators: · p < 0.1, * p < 0.05, ** p < 0.01, *** p < 0.001. **d-f**, Bubble plots showing gene set enrichment analysis (GO Biological Processes and Reactome pathways) for significantly downregulated modules brown and magenta, and significantly upregulated, green module. Chord diagrams displaying the top 5 hub genes (bold) and their highest-weight connections, scaled by Topological Overlap Measure (TOM).

While global ALS synaptic proteomes did not completely segregate from controls on unsupervised principal component analysis (PCA) or hierarchical clustering (**Extended Data Fig. 4a-c**), median-intensity normalization revealed an asymmetric expression signature comprising 487 upregulated and 304 downregulated proteins in ALS (**Extended Data Fig. 4d**), some of which we validated via immunoblotting (**Extended Data Fig. 3**). Stratification by *C9orf72* expansion status revealed localized proteomic variations but did not show the wholesale postsynaptic remodeling we previously documented in end-stage frontal networks^3^, reinforcing the potential capture of early-stage changes in BA17 (**Extended Data Fig. 4e-h**).

To assess the network architecture of this disease state in an unbiased manner, we used Weighted Gene Co-expression Network Analysis (WGCNA) (**Fig. 2a, Extended Data Fig. 5**). Module-trait correlation matrices identified 17 distinct co-expression networks that were significantly associated with the clinical ALS phenotype (**Fig. 2b,c, Extended Data Fig. 5**). Interrogation of the most significantly downregulated network, the *brown* module, revealed a depletion of synaptic vesicle cycle proteins and vesicle fusion machinery via Gene Ontology (GO) and Reactome pathway mapping (**Fig. 2d**). Module eigengene trajectory showed minimal influence of *C9orf72* status (**Extended Data Fig. 6a,c**) and there was a strong correlation between module membership and trait significance (**Extended Data Fig. 6b**). Topological Overlap Measure (TOM) weighting identified the core hub proteins driving this network collapse, isolating essential presynaptic proteins including Synaptogyrin-1 (*SYNGR1*), VGLUT1 (*SLC17A7*), and the essential vesicular proton pump subunits *ATP6V1C1* and *ATP6V1H* (**Fig. 2d**). Importantly, the *magenta* module was also significantly downregulated in ALS and similarly enriched for synaptic vesicle transport terms (**Fig. 2e**). TOM hub analysis identified Synaptophysin (*SYP*) as a principal driver of this network, providing a molecular link to the physical puncta shrinkage captured by our array tomography experiments.

In contrast to this presynaptic vesicular phenotype, the significantly upregulated *green* module was enriched for ribosomal structural proteins, translation initiation, and elongation factors (**Fig. 2f, Extended Data Fig. 6g**). Module eigengene trajectory revealed a greater increase in C9+ve cases (**Extended Data Fig. 6d,f**) and there was a strong correlation between module membership and trait significance (**Extended Data Fig. 6e**). TOM network mapping of the *green* module revealed clusters of large and small ribosomal subunits, which combined with the enrichment of translation initiation and elongation factors, highlights a robust local translational response. Several modules displayed C9-specific profiles, but generally these effects were mild and in smaller modules (**Extended Data Fig. 7**).

Collectively, this unbiased network analysis demonstrates that early-stage visual cortex pathology is characterised by a significant reduction of presynaptic vesicle transport machinery occurring alongside an upregulation of local translation complexes.

### Regional and Disease-Specific Signatures of Visual Cortex Synaptic Proteomes

To determine whether this synaptic vesicle breakdown is a universal, brain-wide feature of ALS, we performed a multi-region multiplexed Tandem Mass Tag (TMT) proteomic analysis (**Fig. 3a**). We pooled BA17 fractions from our donors and compared data directly against our previously published paired primary motor (BA4) and frontal (BA9) cortices from the same individuals. While 4,572 synaptic proteins were conserved across all three regions (**Fig. 3b,c**), BA17 unexpectedly exhibited a significantly higher number of differentially expressed proteins than either BA4 or BA9 (**Fig. 3d**).

**Figure 3.**
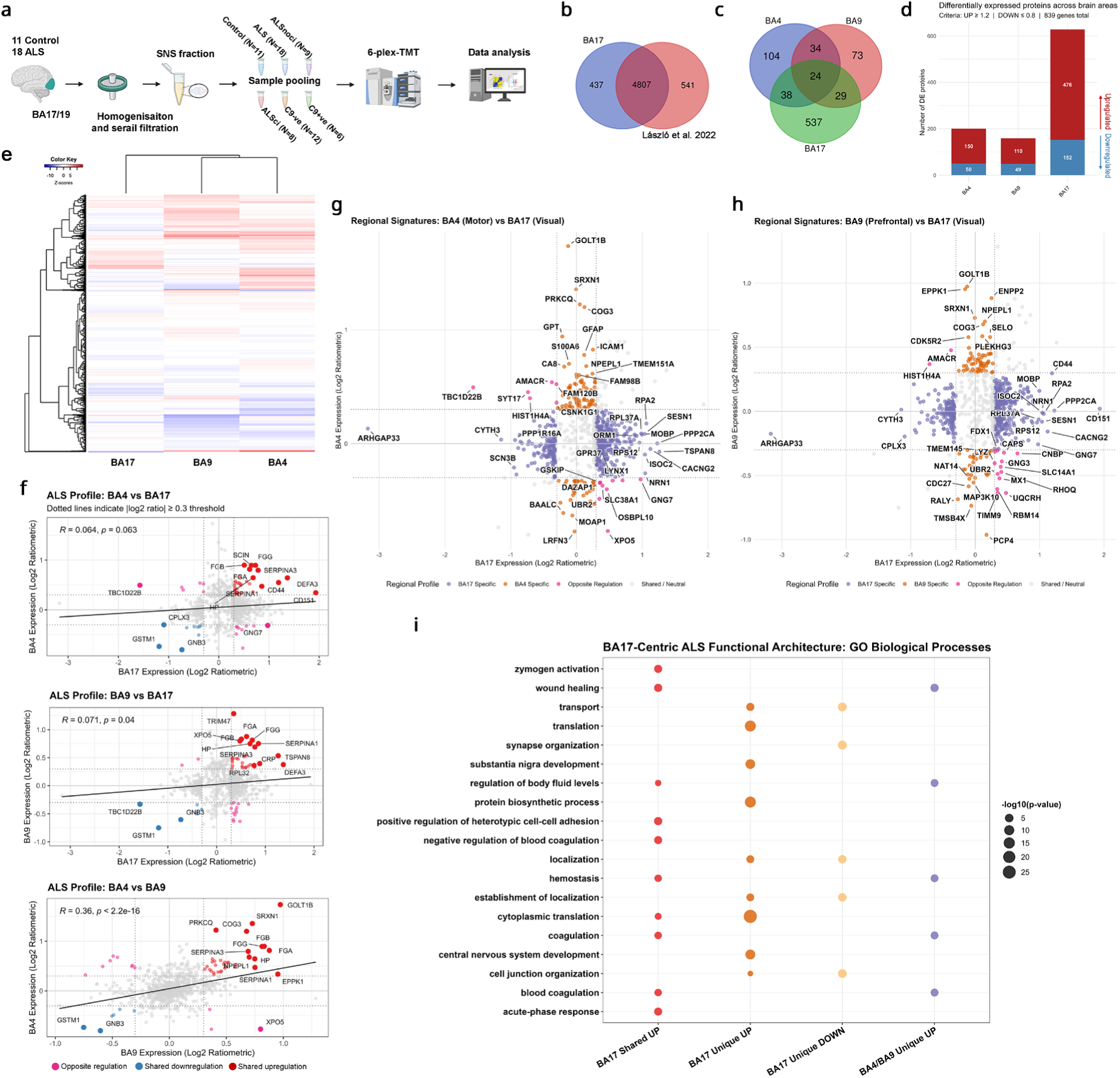
Synaptic proteome mapping across motor, prefrontal, and visual cortices in ALS. **a**, Experimental workflow schematic for synaptoneurosome (SNS) isolation and multiplexed tandem mass tag (TMT) proteomic profiling. Samples were pooled into six disease groups based on cognitive status and *C9orf72* genotype. **b**, Venn diagram illustrating the overlap of 4,572 proteins identified across BA17 and our previous cortical BA4 and BA9 TMT dataset (Laszlo et al. 2022). **c**, Venn diagram showing the overlap of region-specific differentially expressed (DE) proteins, across all three brain areas. **d**, Bar graphs indicating the number of upregulated and downregulated proteins per group (thresholds: ≥1.2-fold upregulated; ≤0.8-fold downregulated). **e**, Unsupervised hierarchical clustering (Euclidean distance) displaying the separation of region-specific proteomic profiles. **f**, Regional cross-comparison of proteomic datasets using pairwise Pearson correlation coefficients (r). **g,h**, Scatter plots of region-specific DE proteins between BA17 and BA4 (**g**) or BA17 and BA9 (**h**). **i**, Bubble plot showing Gene Set Enrichment Analysis (GSEA) of cross-regional TMT data based on region-specific DE proteins.

Unsupervised hierarchical clustering revealed that while the more heavily degenerated BA4 and BA9 networks branched together, BA17 segregated onto its own distinct limb (**Fig. 3e**). Direct ratiometric cross-comparison of log2-transformed data using a conservative 20% biological threshold confirmed this molecular dissociation, showing a tight correlation between BA4 and BA9, but little linear relationship between BA17 and either frontal or motor regions (**Fig. 3f-h**).

Functional pathway mapping using GO bubble plots highlighted the basis of this divergence: while all cortical regions shared a neuroinflammatory profile, BA17 displayed a downregulation in “synapse organization” terms paired with a distinct upregulation in protein translation machinery (**Fig. 3i**).

To determine if presynaptic vesicular protein loss represents an ALS-specific pathology or a generic feature of late-affected cortical breakdown, we aligned our ALS BA17 dataset against our previously published BA17 dataset from Alzheimer’s disease (AD) donors^23^ (**Fig. 4a**). Interestingly, hierarchical clustering separated the cohorts by disease etiology (**Fig. 4c**), and the *ApoE4* positive AD cohort had the highest number of differently expressed proteins (**Fig. 4b,d**). Direct cross-disease analysis between the two most pathologically aggressive genetic subgroups (*C9orf72*-positive ALS and *ApoE4*-positive AD) revealed a shallow correlation, punctuated by cohorts of unique and diametrically opposed protein trajectories (**Fig. 4e,f**).

**Figure 4.**
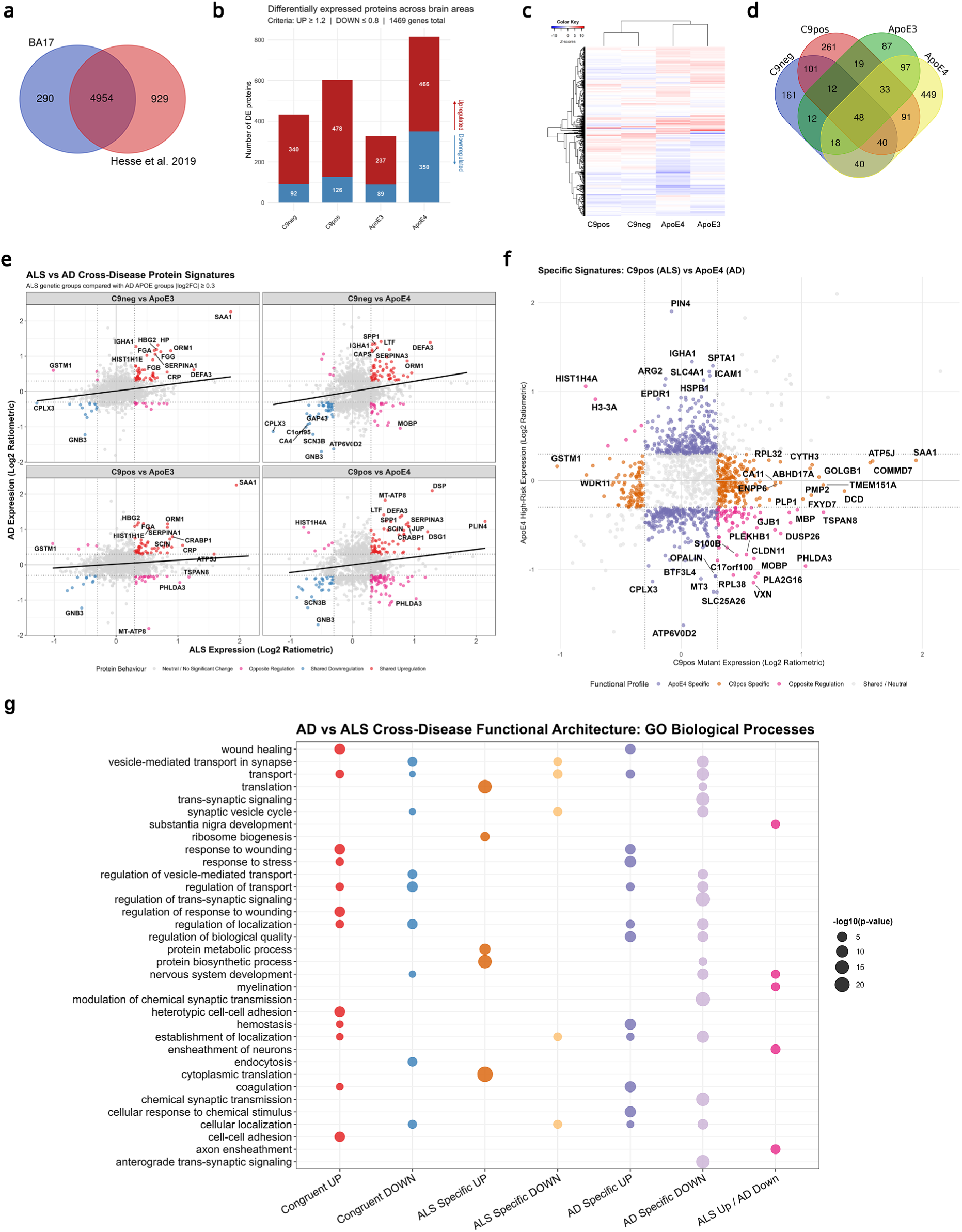
Disease subtype-specific proteomic alterations reveal shared neurodegenerative mechanisms. **a**, Venn diagram showing the overlap of 4,643 proteins between the ALS BA17 TMT dataset and our previous Alzheimer’s disease BA17 TMT proteomic dataset (Hesse et al., 2019). **b**, Bar graphs highlighting disease and genotype-specific protein expression changes. There were 1,469 congruent DE proteins across groups. **c**, Hierarchical clustering (Euclidean distance) demonstrating disease-specific separation among experimental groups. **d**, Venn diagram illustrating disease-specific and overlapping DE protein numbers. **e**, Cross-disease comparison using pairwise Pearson correlation (r) between selected clinical groups. **f**, Scatter plots of disease- and mutation-specific DE proteins between *APOE* ε4 and *C9orf72*-positive groups. **g**, Bubble plot showing GSEA profiles of cross-disease TMT data using disease-specific DE proteins.

GO bubble profiling revealed the upregulation of neuroinflammatory and immune clearance pathways was a shared feature, likely representing a universal tissue response to proteinopathy (**Fig. 4g**), however activation of local translation complexes was ALS-specific. Intriguingly, both ALS and AD cohorts exhibited a significant enrichment of decreased proteins within the ‘synaptic vesicle cycle’ pathway (**Fig. 4g**). However, interrogation at the individual protein level revealed a molecular divergence showing that the proteins differ by disease. This demonstrates that while neurodegenerative diseases converge on the breakdown of the same critical synaptic pathways, they arrive there via disease-specific molecular routes.

### Unbiased Proteomic Alignment Tracks Kinetics of Synaptic Breakdown in Vivo

To resolve the temporal kinetics of this presynaptic breakdown and test its long-term structural viability, we turned to the doxycycline-repressible rNLS8 mouse model of TDP-43 proteinopathy (**Fig. 5a**). Following doxycycline removal, human TDP-43ΔNLS is selectively overexpressed in neurons, driving progressive neurodegeneration. We isolated cortical synaptoneurosomes across five disease timepoints (Weeks 1, 2, 4, 6, and a 6wk + 2-week recovery phase) and analyzed them via label-free DIA proteomics identically to our human cohorts (**Extended Data Fig. 8a-c**).

**Figure 5.**
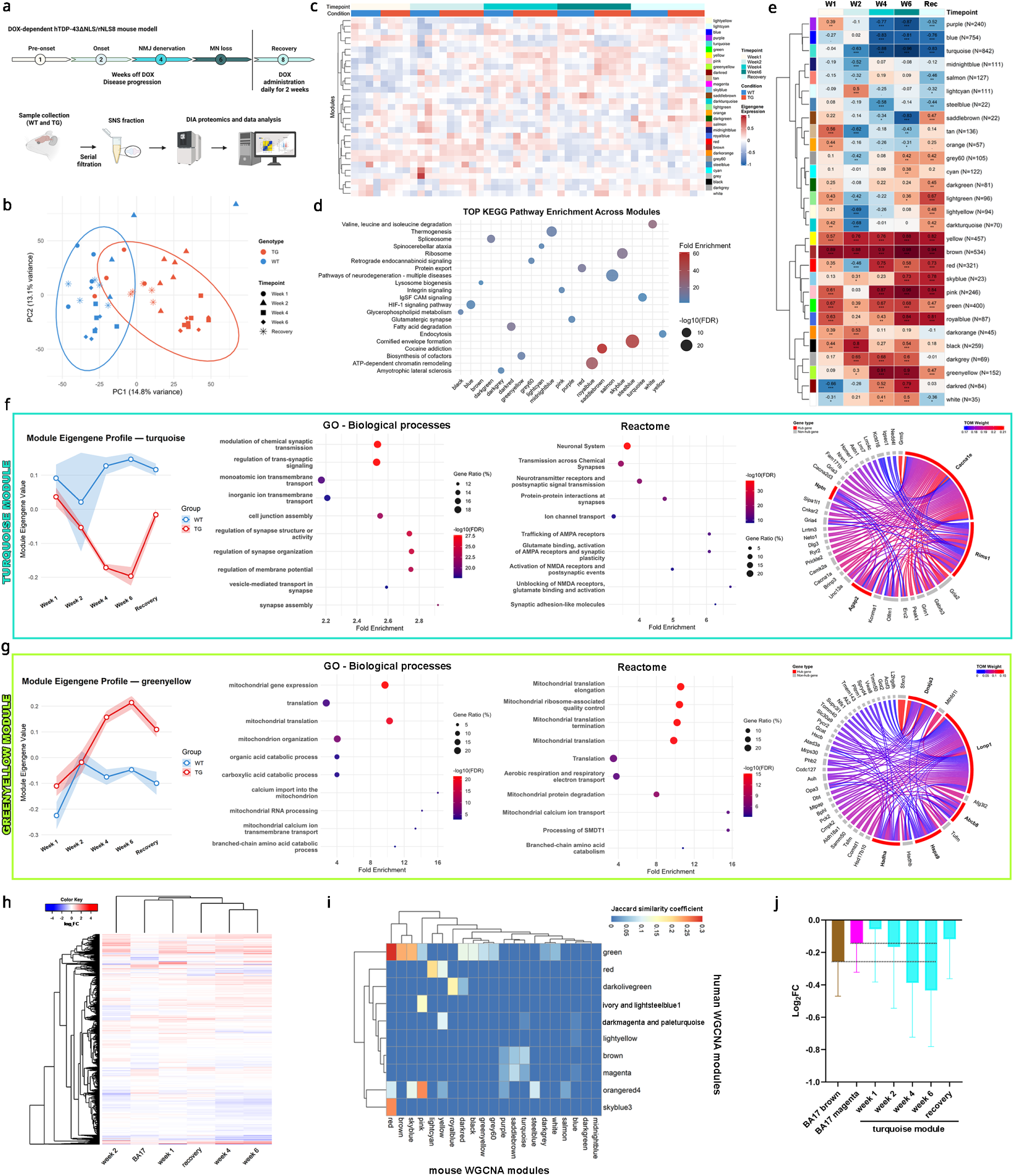
Early synaptic vesicle cycle dysfunction in the hTDP-43ΔNLS mouse model. **a**, Schematic diagram of the experimental and behavioral workflow. **b**, Principal component analysis (PCA) of the synaptic proteome from hTDP-43 rNLS8 mouse brain samples. Colors indicate genotypes; shapes indicate longitudinal collection timepoints. **c**, Adjacency heatmap showing hierarchical clustering of module eigengenes per sample to illustrate inter-module relationships based on eigengene correlation. **d**, Bubble plot displaying the most significantly enriched KEGG pathways for trait-associated modules. **e**, Hierarchical heatmap of significant module–trait correlations from weighted gene co-expression network analysis (WGCNA) using biweight midcorrelation (bicor). Significance levels: · p < 0.1, * p < 0.05, ** p < 0.01, *** p < 0.001. W1–W6, weeks 1–6; Rec, recovery stage. **f,g**, Trajectory plots of sample–module relationships for the turquoise and greenyellow modules (lines indicate mean ± s.d.) accompanied by functional enrichment profiles (GO Biological Processes and Reactome pathways) and chord diagrams displaying the top 5 hub genes and their highest-weight connections scaled by Topological Overlap Measure (TOM). **h**, Hierarchical clustering (Euclidean distance) of the log2 fold changes comparing human BA17 DIA data and the rNLS8 mouse model disease stages. **i**, Heatmap of pairwise Jaccard similarity coefficients comparing GO Biological Process terms derived from human and mouse WGCNA modules. Rows and columns are ordered by hierarchical clustering using Euclidean distance. **j**, Bar graphs showing log2 fold changes for downregulated synaptic modules across species (human: brown and magenta modules; mouse: turquoise module). Genotypes: WT, wild-type; TG, transgenic animals overexpressing hTDP-43ΔNLS. Number of animals was used: Week1 WT=4, TG=4, Week2 WT=4, TG=6, Week4 WT=4, TG=6, Week6 WT=5, TG=5, Recovery WT=3, TG=3.

Synaptic alterations occurred as early as 1-week post-induction (pre-disease onset stage), escalating continuously through week 6 (late-stage disease) (**Extended Data Fig. 8d-g,i**). Remarkably, re-introducing doxycycline for just 2 weeks at week 6 drove the global late-stage synaptic proteome backwards in time, causing it to resemble the milder, early disease states (**Extended Data Fig. 8h,i**). Unsupervised PCA largely separated the cohorts by genotype and chronological disease stage (**Fig. 5b**).

WGCNA network mapping of the longitudinal mouse dataset identified discrete co-expression modules that tracked with disease progression (**Fig. 5c-e, Extended Data Fig. 9**). The *turquoise* module exhibited a continuous decline from week 1 through week 6, followed by a dramatic upward reset back close to control levels during the recovery window (**Fig. 5f**). GO and Reactome mapping of this module isolated essential parameters of synaptic transmission, highlighting key vesicle primers (*RIMS1, UNC13A*) and voltage-gated calcium channel subunits (Cav2.3) (**Fig. 5f**). *Purple* and *saddlebrown* modules also longitudinally decreased, and these were enriched for other essential vesicular proteins including synaptotagmin-1 (*SYT1*), synaptic vesicle proteins 2A/2B *(SV2A/B*) and vesicular monoamine transporter 2 (*SLC18A2*) (**Extended Data Fig. 10**).

In striking contrast to the progressive decline observed in the *turquoise* network, the *greenyellow* module **(Fig. 5g)** exhibited a mirrored, inverse trajectory. Eigengene protein mapping revealed a continuous increase in expression from Week 1 through Week 6, which partially reversed back towards baseline during the 2-week recovery stage. Functional profiling of this module via GO and Reactome analysis revealed enrichment for mitochondrial oxidative phosphorylation, ribosomal biogenesis, and cellular stress response pathways **(Fig. 5g)**. *Brown* and *yellow* modules also displayed progressive increases in expression, and these were enriched for pathways involved in autophagy, ER stress, and protein misfolding (**Extended Data Fig. 10**).

To map our human postmortem findings onto this *in vivo* timeline, we performed cross-species hierarchical clustering. Strikingly, the human BA17 synaptic proteome clustered alongside Week 1 mouse stage, providing strong evidence that the human visual cortex displays an early, pre-symptomatic window of ALS pathology (**Fig. 5h**). Jaccard similarity matrices confirmed that this cross-species match was driven by an overlapping molecular signature: the downregulated human *brown* and *magenta* networks exhibited significant overlap with the mouse *turquoise* and *purple* vesicle modules (**Fig. 5i, Extended Data Fig. 11**). Finally, expression-scale distribution alignment of these modules positioned the human presynaptic deficit between weeks 2 and 4 of the mouse timeline, highlighting the transition period from pre-disease onset to early stage disease^19^ (**Fig. 5j**).

### Rebalance of Synaptic Change Following Removal of Pathology

In the final phase of our study, we exploited the rNLS8 model’s inducible nature to define the molecular mechanics of circuit recovery. We isolated proteins that exhibited a significant disease-associated trajectory (upregulation or downregulation) followed by a return (complete or partial) to control baselines upon transgene repression (**Fig. 6a-c**). Unsupervised clustering confirmed that the recovery stage clustered with early pre-symptomatic profiles, suggesting a rebalance of the synaptic proteome (**Fig. 6a**).

**Figure 6.**
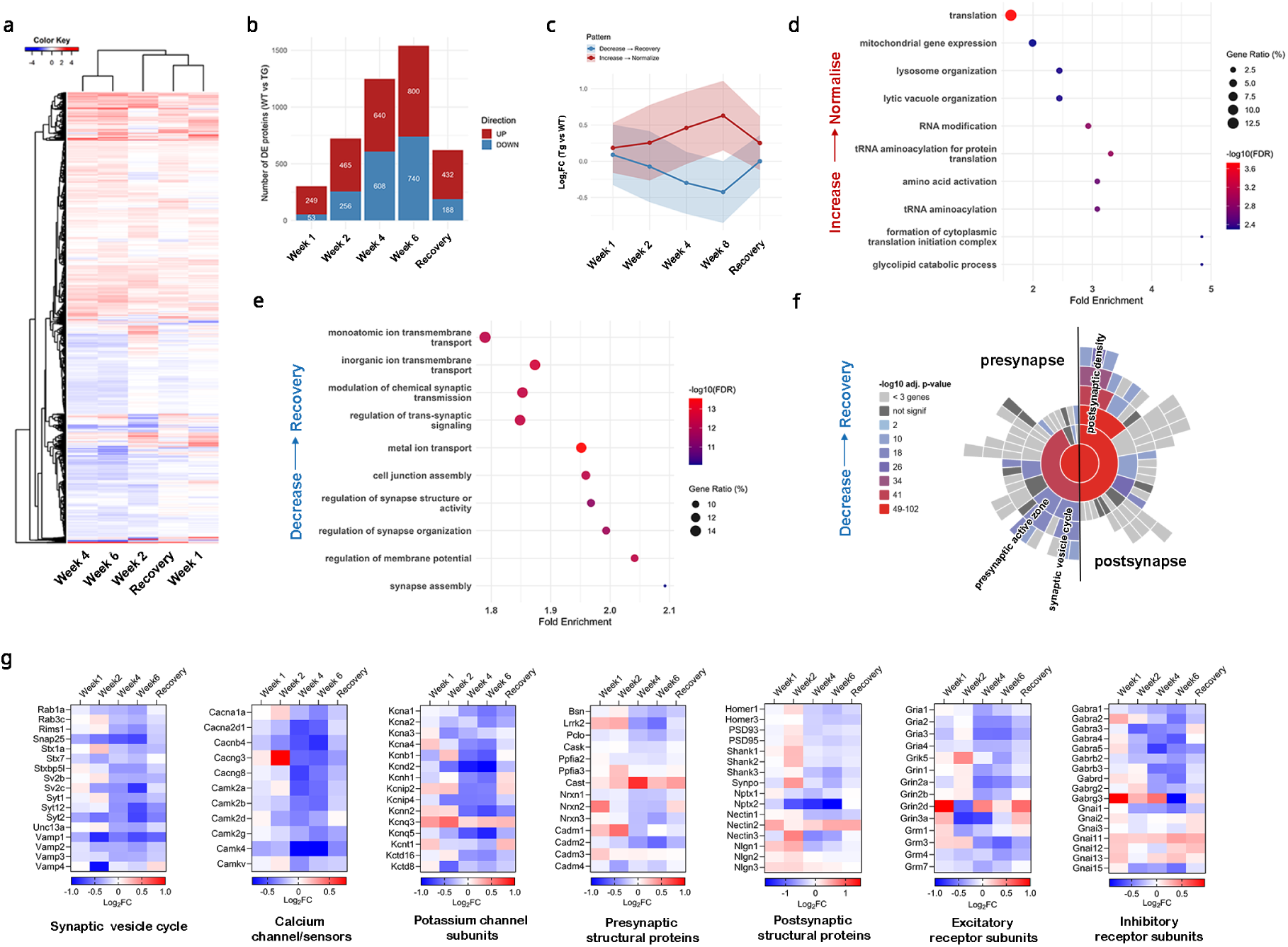
Significant synaptic proteome recovery following TDP-43 pathology clearance. **a**, Hierarchical clustering (Euclidean distance) showing proteomic separation between longitudinal disease and recovery stages. **b**, Bar graphs showing the number of DE proteins identified per timepoint (n = 1,948 total DE proteins across all phases). **c**, Longitudinal expression trajectories of DE proteins clustered into decreased-and-recovered (n = 1,117) or increased-and-recovered (n = 831) expression profiles. **d-e**, Bubble plots showing GSEA functional profiles (GO Biological Processes) across the distinct expression trajectories. **f**, Sunburst plot displaying pre- and postsynaptic cellular component enrichment within the decreased-and-recovered trajectory. **g**, Heatmaps illustrating the relative abundance patterns of key synaptic proteins across progressive disease phases and following the recovery period.

Network analysis of this salvageable proteome revealed a reciprocal rebalancing: the activated local translation and ribosomal processes (up-and-recover) were deactivated, while the networks governing synapse assembly, membrane potential regulation, and trans-synaptic signaling (down-and-recover) were restored (**Fig. 6d,e**). Subsynaptic localization mapping reinforced that this rescue involved changes within the presynaptic vesicle compartments, amongst others including postsynaptic signalling complexes (**Fig. 6f**).

Targeted heatmaps traced recovery within groups of calcium-sensitive sensors, regulatory potassium channels that control terminal excitability, and large cascades of pre- and post-synaptic signaling proteins which all displayed a synchronized, comprehensive restocking of their protein levels back to homeostatic baselines (**Fig. 6g**).

Crucially, to determine if remaining terminals are structurally viable during prolonged presynaptic breakdown, we tracked the longitudinal trajectories of core trans-synaptic structural frameworks within the mouse fractions. While vesicle proteins progressively drop from week 2, the primary trans-synaptic adhesion complexes, including Neurexin-1 (*NRXN1*), Synaptic Cell Adhesion Molecule 1 (*CADM1*), and the core presynaptic active zone plasma membrane anchor (*CASK*), remained stable across all disease weeks, as did the postsynaptic scaffolding elements *SHANK1 and SHANK2* (**Fig. 6g**). In contrast, vesicle-docking scaffolds (*Bassoon, Piccolo*) and functional receptor-clustering complexes (*DLG4, NLGN1*) remained stable early on but followed the hub proteins into a delayed decline at week 6. Importantly, except for NRXN1(which decreases), all these proteins were also unchanged in human ALS BA17.

Because the trans-synaptic cell-adhesion structures and active zone anchors are preserved throughout the early disease window, the surviving terminals do not need to be structurally rebuilt *ex novo*. Instead, they are available to restock vesicle machinery once pathogenic TDP-43 expression is repressed. Collectively, these data demonstrate that early synaptic alterations in ALS represent a structurally salvageable state of neurodegeneration.

## Discussion

Here, we identify the primary visual cortex (BA17/19) as a unique anatomical region in which the earliest human cortical changes associated with ALS can be examined before extensive neuronal degeneration occurs. The dissociation between cortical thinning and preserved neuronal density argues against the common assumption that regional atrophy directly reflects neuronal loss. Instead, our findings indicate that the earliest structural change is a reduction of the neuropil compartment while neuronal cell bodies remain largely preserved. This observation implicates synaptic and axonal compartments, rather than neuronal somata, as the primary sites of early injury, consistent with the concept of dying-forward neurodegeneration^24,25^.

Our data further suggest that this cortical compaction may be driven by degeneration of afferent white matter pathways rather than an intrinsic cortical process. Ultrastructural axonal pathology, widespread myelin fragmentation, nuclear loss and astrogliosis in the underlying white matter of the visual cortex occurred alongside an absence of pronounced reactive gliosis within the overlying cortex, revealing a striking dissociation between grey and white matter pathology. Together, these observations suggest that the cortical grey matter initially responds to degenerating afferent projections rather than suffering intrinsic pathological mechanisms^26,27^. Notably, phosphorylated TDP-43 pathology (although mild) was detected in the majority of BA17 samples, especially when present in earlier-affected motor and frontal cortices, supporting the theory that pathological TDP-43 spreads through anatomically connected neuronal networks in a temporal manner, potentially contributing to early axonal dysfunction and synaptic failure^11,18^.

Using complementary high-resolution and ultrastructural imaging approaches, we found that remaining synapses in BA17 undergo morphological remodelling. Presynaptic terminals exhibited reduced volume and accumulation of electron-dense material indicating that terminals are proteostatically stressed and shrinking. These alterations matched our proteomic findings, which demonstrated depletion of presynaptic proteins involved in vesicle trafficking while trans-synaptic scaffold proteins were preserved.

Consistent with these observations, WGCNA identified disease-associated modules that were highly enriched for components of the synaptic vesicle cycle. Hub proteins included regulators of vesicle identity (SYP, SYNGR1), glutamate loading (VGLUT1) and vesicle acidification (ATP6V1C1 and ATP6V1H), indicating that the earliest molecular defects affect neurotransmitter release rather than global synaptic architecture. These findings support the concept that functional synaptic impairment precedes structural synapse loss^8^.

Our proteomics data also revealed downregulation of calcium-binding proteins and ion channel regulators, including NECAB2 and CAMKIIA, which is predicted to alter calcium homeostasis, receptor trafficking and ion channel function at both pre- and postsynaptic compartments. Furthermore, our longitudinal analysis of the rNLS8 mouse model demonstrated that multiple calcium and potassium channel components were already reduced one week after induction of human ΔNLS-TDP-43 expression. Reduced potassium conductance would impair action potential repolarisation, promoting repetitive firing and excessive glutamate release, while disrupted calcium buffering further compromises calcium-dependent potassium channel activity^28–30^. Together, these changes provide a plausible mechanism linking early ion channel dysfunction to neuronal hyperexcitability, a phenomenon widely recognised as an early feature of ALS^31,32^.

Calcium signalling also plays a central role in synaptic vesicle cycling. CaMKII regulates synapsin I phosphorylation, controlling mobilisation of reserve vesicles, whereas calcium-sensitive priming proteins such as Munc13 or the ALS-related UNC13A facilitate SNARE complex assembly and vesicle fusion^33,34^. Reduced calcium influx together with impaired calcium sensing would therefore be expected to compromise vesicle recruitment, priming and neurotransmitter release^35^. Our findings suggest that disruption of calcium signalling represents an upstream event contributing to the selective impairment of the synaptic vesicle cycle observed in both human tissue and the rNLS8 model. Importantly, the calcium channel blocker nifedipine is currently being investigated in the UK-wide EXPERT-ALS clinical trial.

The rapid onset of these molecular changes following neuronal expression of ΔNLS-TDP-43 further suggests that TDP-43 proteinopathy initially disrupts synaptic function through selective impairment of ion channel regulation and vesicle cycling rather than widespread synaptic disassembly. This may reflect loss of TDP-43-mediated regulation of transcripts encoding presynaptic proteins such as the well-described ALS targets UNC13A and KCNQ2, or mislocalisation of TDP-43-dependent RNA-binding complexes required to maintain local synaptic protein homeostasis.

In contrast to the decline in vesicle-associated proteins, WGCNA identified a reciprocal increase in ribosomal proteins and translation-initiation machinery. We interpret this response as an adaptive attempt to preserve local protein synthesis rather than as a parallel pathogenic pathway. However, recent work also suggests that increased abundance of orphan ribosomal proteins may reflect accumulation of unassembled ribosomal subunits rather than increased translational output^36^. Importantly, our translational signature also contained initiation and elongation factors, and closely paralleled the decline in vesicle proteins in the longitudinal mouse dataset and reverted following suppression of ΔNLS-TDP-43 expression together with restoration of the vesicle proteome. These observations support the interpretation that presynaptic terminals are attempting to compensate for loss of vesicle-associated proteins by enhancing local translational capacity. Whether this response is ultimately protective, insufficient or maladaptive during disease progression remains an important question for future investigation.

Although the motor cortex is the principal site of pathology in ALS, increasing evidence indicates that multiple cortical regions are affected during disease progression, and that this involvement cannot be explained solely by the distribution of TDP-43 pathology. While approximately 90% of ALS cases exhibit phosphorylated TDP-43 inclusions, cortical dysfunction can also occur in patients with little or no detectable TDP-43 pathology, suggesting that disease-relevant circuit dysfunction may arise independently of overt protein aggregation^2^. By comparing the primary motor cortex (BA4), dorsolateral prefrontal cortex (BA9) and primary visual cortex (BA17) from the same individuals, we found that BA17, despite representing the least pathologically advanced region, displayed the greatest number of differentially expressed proteins and formed a distinct molecular cluster in unsupervised analyses. Such differences may arise from intrinsic properties of the visual cortex, including its unique laminar organisation, neuronal activity and cellular composition, alternatively, ALS-driven synaptic breakdown is the same across the CNS and BA17 is at an earlier stage on the pathological timeline than BA4 and BA9.

Comparison with Alzheimer’s disease further reinforces the distinction between pathway-level convergence and molecular specificity. Despite their different initiating pathologies^23^, both ALS and Alzheimer’s disease showed disruption of the synaptic vesicle cycle together with activation of neuroinflammatory and immune-related pathways, suggesting that these processes represent common responses to cortical neurodegeneration^23,37,38^. However, the individual proteins underpinning these shared pathways differed substantially between diseases, and the increased abundance of ribosomal and translational machinery observed in ALS was not evident in Alzheimer’s disease. Thus, while pathway enrichment identifies common biological processes affected across neurodegenerative disorders, it does not necessarily imply conservation of the underlying molecular mechanisms. This distinction has important implications for therapeutic development, as interventions directed against individual components of a shared pathway in one disease may not be directly applicable to another. Our findings therefore emphasise the importance of disease-specific proteomic profiling alongside pathway-level analyses to identify molecular targets that are both biologically relevant and therapeutically tractable in ALS.

Cross-species integration placed the molecular profile of human BA17 between the early pre-disease onset and early disease stages of the rNLS8 mouse model, preceding the widespread synaptic dysfunction observed at six weeks after induction. Independent WGCNA and longitudinal trajectory analyses demonstrated conservation of disease-associated molecular programmes across species. These included coordinated disruption of presynaptic vesicle cycling, calcium-dependent neurotransmission, axonal cargo transport, and synaptic maintenance pathways, with reductions in key components involved in vesicle docking and recycling (e.g., SNARE-associated proteins, RAB GTPases, and synaptic vesicle regulators), calcium-dependent signalling (a wide range of calcium and potassium channel subunits), and neuronal transport machinery (KIF5A, KIF17, KIF21B). In parallel, activation of translational pathways preceded overt structural degeneration, potentially suggesting an early compensatory response aimed at maintaining synaptic protein homeostasis under conditions of increased proteostatic stress. The preservation of these conserved molecular signatures across human and mouse tissue supports the utility of the rNLS8 model for investigating early synaptic dysfunction in ALS, while recognising that acute transgene-driven ΔNLS-TDP-43 expression does not fully capture the prolonged, heterogeneous, and multifactorial nature of sporadic disease.

An important observation from the mouse model is that these early molecular alterations were largely reversible following suppression of ΔNLS-TDP-43 expression. Recovery of synaptic vesicle-associated proteins occurred alongside normalisation of translational programmes, indicating that presynaptic terminals retain the capacity to restore molecular homeostasis before progression to irreversible structural degeneration^22^. This is consistent with findings demonstrating recovery of behavioural and motor phenotypes following a longer six-week recovery period in the same model^39^. Together, these observations suggest that early synaptic dysfunction represents a dynamic and potentially modifiable phase of disease progression. However, translating this concept to patients will require the development of biomarkers capable of identifying this early window *in vivo*, as well as therapeutic strategies that intervene before synaptic destabilisation progresses to permanent circuit disruption.

Together, our findings define an early stage of cortical pathology characterised by selective impairment of synaptic function, particularly involving presynaptic vesicle regulation, neurotransmitter release machinery, calcium homeostasis, and neuronal transport, while the broader synaptic architecture remains relatively preserved. The accompanying increase in translational activity may represent a transient adaptive response to maintain presynaptic protein supply and compensate for early synaptic stress. Rather than representing an irreversible process from disease onset, these data support a model in which molecular synaptic dysfunction precedes structural synapse loss and neuronal degeneration, thereby identifying an actionable therapeutic window. Future studies should focus on establishing biomarkers that detect this stage in living patients and developing interventions aimed at preserving or restoring presynaptic function before irreversible synaptic elimination occurs.

## Methods

### Donor characteristics and brain collection

Fresh frozen human tissue was collected and handled as described previously^3^. All ALS donors were clinically characterised according to the revised El Escorial criteria and recruited via the Scottish Motor Neurone Disease Register, with clinical, genetic, and cognitive data obtained from the CARE-MND database. Cognitive status was assessed using the Edinburgh Cognitive and Behavioural ALS Screen (ECAS), and genetic profiling was performed as previously described^40^. All procedures involving human participants and post-mortem tissue were approved by the relevant Scottish ethics committees, including the Edinburgh Brain Bank and ACCORD. Post-mortem tissue from the primary visual cortex (Broadmann area 17/19), was obtained from clinically diagnosed ALS cases (n = 11) and age- and sex-matched neurologically normal controls (n = 18, Supplementary Table 1). Detailed demographics can be found in Additional file 1.

**Supplementary Table 1.** Summary demographics of all donors.

|  | Number of cases | Male/Female | Post-mortem interval (hrs, median+range) | Age at death (years, median+range) | Genetics | Region of onset |
| --- | --- | --- | --- | --- | --- | --- |
| **Cohort 1 – Electron microscopy** |  |  |  |  |  |  |
| Control | 6 | 4/2 | 57 (40-99) | 58.5 (53-77) |  |  |
| ALS | 20 | 11/9 | 83 (35-131) | 62.5 (40-89) | 5 - c9orf72+ve | 5 bulbar, 12 limb, 1 mixed |

Supplementary Table 1. Summary demographics of all donors.
| Cohort 2 - Neuropathology |  |  |  |  |  |  |
| --- | --- | --- | --- | --- | --- | --- |
| Control | 11 | 7/4 | 68 (39-77) | 79 (77-82) |  |  |
| ALS | 23 | 15/8 | 77 (30-131) | 62 (43-83) | 6 - c9orf72+ve<br>2 - NEK1+ve<br>2 - SOD1+ve | 7 bulbar,<br>13 limb, 2 mixed |
| Cohort 3 - Proteomics |  |  |  |  |  |  |
| Control | 11 | 6/5 | 74 (3-115) | 71 (57-85) |  |  |
| ALS | 23 | 10/13 | 86.5 (30-131) | 62 (43-83) | 6 - c9orf72+ve | 6 bulbar,<br>11 limb, 1 mixed |

### Tissue collection of hTDP-43- ΔNLS mouse model

Mice used for proteomic analysis were obtained by intercrossing homozygous tetO-hTDP-43-ΔNLS line 4 (RRID:IMSR_JAX:014650) mice with hemizygous NEFH-tTA line 8 (RRID:IMSR_JAX:025397) mice ^19^, on a pure C57Bl/6JAusb background at The University of Queensland, as previously described^22,39^. Single transgenic hemizygous tetO-hTDP-43-ΔNLS line 4 littermates were used as controls, and genotyping was performed using tail DNA. Experiments were conducted with approval from the Animal Ethics Committee of The University of Queensland (#QBI/131/18 and 2022/AE000578). Mice were group housed in a Specified Pathogen-Free (SPF) animal facility, and were housed with a 12-hour light/dark cycle (lights on beginning at 6am). Room temperature and humidity were kept constant at 21 ± 1°C and 55 ± 5%, respectively. All mice were fed Doxycycline-containing chow (200 mg/kg) until a mean age of approximately 10 weeks, when food was replaced with standard chow lacking Doxycycline – this facilitates expression of the hTDP43-ΔNLS transgene. Following removal of Doxycycline, mice were scored 3 times per week against a neurological monitoring sheet, to confirm disease onset and monitor progression. Disease onset was defined when hindlimb clasping was reported for 2 consecutive sessions. Doxycycline was reintroduced into the diet 6 weeks later, and mice were assessed for a further 2 weeks to monitor the recovery phase (Supplementary Table 2).

**Supplementary Table 2.** Details of the hTDP-43/rNLS8 mice were used in the study.

| Timepoint | Genotype | N + Sex |
| --- | --- | --- |
| Week 1 | Littermate control ('WT') | 4 (2 male + 2 female) |
|  | Transgenic rNLS8 | 4 (2 male + 2 female) |
| Week 2 | Littermate control ('WT') | 4 (2 male + 2 female) |
|  | Transgenic rNLS8 | 6 (2 male + 4 female) |
| Week 4 | Littermate control ('WT') | 4 (1 male + 3 female) |
|  | Transgenic rNLS8 | 6 (3 male + 3 female) |
| Week 6 | Littermate control ('WT') | 5 (1 male + 4 female) |
|  | Transgenic rNLS8 | 5 (1 male + 4 female) |
| Recovery | Littermate control ('WT') | 5 (3 male + 2 female) |
|  | Transgenic rNLS8 | 5 (3 male + 2 female) |

### Neuropathology

Immunohistochemistry was performed as described previously^2^. Briefly, fresh post-mortem tissue blocks were fixed in 10% formalin, processed for paraffin embedding, and sectioned at 4 μm using a Leica microtome. Sections were stained using standard immunohistochemistry protocols with the Novolink Polymer detection system and DAB chromogen, followed by hematoxylin counterstaining. Antibody details are provided in Supplementary Table 3. Appropriate positive and negative controls were included for each staining run, and all analyses were performed blinded to clinical diagnosis. For pTDP-43 and pTau assessment, cases were classified as positive or negative based on the presence or absence of clearly defined protein aggregates.

### QuPath analysis

Post-mortem human samples were sectioned, stained by the Edinburgh Brain Bank, and digitised using whole-slide scanners at the University of Glasgow and the University of Dundee (NanoZoomer S60 Digital Slide Scanner – 40 X objective, Zeiss Axio Scan 7 slide scanner – 20X objective). Whole-slide images were analysed using QuPath (version 0.6.0).

Regions of interest (ROIs) corresponding to grey matter and white matter were manually annotated using the freehand selection tool. Image preprocessing included colour deconvolution (H-DAB) to separate haematoxylin and DAB signals. Cell detection was performed by using the Watershed Cell Detection algorithm on the hematoxylin optical density channel for measuring cell density in the white matter and DAB channel for calculating NeuN-positive cell density. For white matter annotations, detection was performed at 0.45 µm pixel size with a smaller background radius (5 µm), reduced smoothing (σ = 1.5 µm), and a lower threshold (0.1); watershed post-processing was enabled, and nuclei were expanded by 5 µm. Boundary smoothing was applied, nuclei were included, and quantitative measurements were generated for all detected cells.

NeuN-positive cell populations were identified using Positive cell detection, and intensity profiles were set based on the suggested autothreshold by Qupath. The amount of NeuN-positive cells was counted and normalised to ROI area to get density results. Classification was validated by manual measurement of nuclei in haematoxylin and eosin (H&E) and NeuN-stained sections.

Immunohistochemical quantification of GFAP and CD68 staining was performed using DAB-based positive pixel detection. Thresholds for DAB positivity (weak, moderate, strong) were defined and applied consistently across all images. Staining burden was quantified as the percentage of DAB-positive area within each ROI.

Cortical grey matter thickness was assessed at ten randomly selected points per section by measuring the distance between the pial surface and the grey–white matter boundary below cortical layer 6. The mean of these measurements was used as the representative cortical thickness for each brain region.

All analyses were performed blindly and visually inspected to ensure accurate segmentation and to exclude artefacts (e.g., tissue folds, staining irregularities).

### Generation of synaptoneurosome samples

Synaptoneurosome samples were prepared as descired earlier^3,5^. Briefly, fresh-frozen human brain tissue from BA17/19 was homogenised on ice in homogenisation buffer (25 mM HEPES, pH 7.5; 120 mM NaCl; 5 mM KCl; 1 mM MgCl₂; 2 mM CaCl₂) supplemented with protease and phosphatase inhibitors. The homogenate was first passed through an 80 µm nylon mesh to generate the total homogenate (TH), from which an aliquot was retained. The remaining homogenate was further filtered through a 5 µm pore-size filter to obtain a synaptoneurosome (SNS)-enriched fraction. This fraction was centrifuged (5 min, 1000 × g), and the resulting pellet was washed twice in homogenisation buffer. The final pellet was weighed and resuspended in 100 mM Tris-HCl (pH 7.6) containing 4% SDS and protease inhibitors (1:5 w/v based on pellet weight), followed by further homogenisation and centrifugation (20 min, 17,000 × g, 4 °C). The supernatant was collected, and protein concentration was determined using a Micro BCA assay. Samples were stored at −80 °C until further experiments.

Samples were individually analysed using Data Independent Acquisition (DIA) Proteomics and 25 µg of protein was also pooled into 6 representative disease groups based on genetics and cognitive performance replicating the experimental setup of Laszlo et al 2022 for 6-plex TMT (Suppl fig X). The 6 pools were as follows: Pool 1—Control (n = 11); Pool 2—ALS (n = 18); Pool 3— ALSnoci (n = 9); Pool 4— ALSci (n = 8); Pool 5—*C9orf72*-ve (n = 12); Pool 6—*C9orf72* +ve (n = 6).

### Western blot

Western Blots were performed using human post-mortem samples as previously described^3^. Briefly, protein concentrations were determined via the Pierce Micro BCA assay (Thermo Fisher, #23235), then 10 µg of each sample was prepared with 4X Laemmlie sample buffer (Bio-Rad, #1,610,747), supplemented with beta-mercaptoethanol. Samples were denatured at 95°C for 5 minutes, then loaded into 4-20% Tris-Glycine 1.0 mm polyacrylamide pre-cast gels (Thermo Fisher, # WXP42020BOX), with 8 µL protein ladder (Li-Cor, #928-70000). Proteins were subsequently transferred onto nitrocellulose membranes, using precast transfer stacks (Invitrogen, #IB23001), with the iBlot™ 2 Gel Transfer Device (Invitrogen, #IB21001). Following transfer, total protein staining was performed using REVERT™ 700 Total Protein stain (Li-Cor, REVERT™ 700 Total Protein Stain Kits, #926–11,010), and membranes were imaged with the Li-Cor Odyssey System. Membranes were destained following the manufacturers instructions, then blocked for 1 hour at room temperature with 5% milk/TBST or 5% BSA/TBST. Membranes were incubated overnight at 4°C in primary antibodies, diluted in the appropriate block solution. After thorough washing in TBST, membranes were incubated for 1 hour at room temperature in secondary antibodies, prepared in blocking solution. Membranes were finally imaged as before.

### Tandem mass tagged (TMT) liquid chromatography mass spectrometry (LC–MS/MS)

6-plex Tandem Mass Tag (TMT) proteomics was performed as previously described^3^. After S-trap processing samples were labelled with TMT tags using Pierce High pH Reversed-Phase Peptide Fractionation kit (Thermo Scientific, #) following the manufacturer’s protocol. TMT tages are the following: 126- Control, 127N- ALS, 128C-ALSnoci, 129N- ALSci, 130C- C9-ve, 131- C9+ve

Mass spectrometry analysis was carried out at the ‘FingerPrints’ Proteomics Facility, Faculty of Life Sciences, University of Dundee. Analysis of peptides was performed on a Q-Exactive-HF (Thermo Scientific) mass spectrometer coupled with a Dionex Ultimate 3000 RSLC Nano (Thermo Scientific) as described before (Laszlo et al 2022). Collected peptide identification was made using MaxQuant (version 1.6.2.10) and searched against the SwissProt subset of the H. sapiens Uniprot database (May 2019 release) using the Andromeda search engine software^3^.

### Data-independent acquisition (DIA) proteomics and mass spectrometry

Protein quantification was performed using a MicroBCA kit (Pierce). Samples were processed using S-Trap Mini columns (Protifi) where proteins were reduced with dithiothreitol (VWR proteomics grade), alkylated with iodoacetamide (Sigma) and digested overnight with 2.75µg of trypsin (Pierce MS-grade) at 37°C. A second digest was repeated the following day for a further 6 hours before peptides were eluted from the columns and dried. Digested peptides were then resuspended in 40 µl 1% formic acid. Resuspended peptides were run on an Orbitrap Astral mass spectrometer (Thermo Scientific) coupled to a Vanquish Neo UHPLC system (Thermo Scientific) with LC buffers compromising of buffer A (0.1% formic acid) and buffer B (80% acetonitrile, 0.1% formic acid). The buffers were used to create a gradient for a 60SPD (samples per day) run where peptides were trapped on a PepMap Neo C18 trap cartridge (Thermo Scentific 174500) before being eluted from a PepMap RSLC C18 column (Thermo Scientific ES906). Peptides were eluted at a flow rate of 800nl/min with a gradient starting at 4%B and increasing to 8%B within 0.6 minutes, from 8%B to 22.5%B in the next 13 minutes, from 22.5%B to 35%B in 6.9 minutes and then increasing to 55%B for the last 0.4 minutes. The column was then washed and equilibrated. The Orbitrap Astral was operated in positive ionisation mode, equipped with an EasySpray source. The source voltage was set to 1.90kV and the ion transfer tube was set to 280°C.

RAW data was acquired in Data Independent Acquisition mode and a scan cycle comprised a full MS (MS1) scan followed by MS/MS DIA scans. For the MS1 scan, data was collected in profile mode with a resolution of 240,000, custom Automatic Gain Control (AGC) target of 300%, maximum injection time (IT) of 5ms, microscan of 1 and a mass range of 380-980m/z was used.

MS/MS DIA scans were performed in centroid mode with window widths of 4 Th and an overlap of 0m/z. HCD collision energy of 25%, a custom AGC target of 500, maximum IT of 3ms and a microscan of 1. The detector type used was Astral with a scan range of 150-2000m/z.

All raw files and proteomics data have been deposited to the ProteomeXchange Consortium via the PRIDE partner repository (https://www.ebi.ac.uk/pride/) with the dataset identifier (*will be updated upon publication*).

### Bioinformatic analyses

#### Data-independent acquisition (DIA) proteomics

Thermo .RAW files were converted to HTRMS files prior to data analysis using HTRMS converter (Biognosys). Data analysis was performed using Spectronaut version 19 (Biognosys) using directDIA. Samples were searched against a mouse database with variable modifications of Oxidation(M), Dioxidation(MW), Acetyl (Protein N-term), Deamidation (NQ) and Gln->Pyro-Glu set and a fixed modification of Carbamidomethyl (C). For identification a precursor PEP (Posterior Error Probability) cutoff of 0.1 and a protein PEP cutoff of 0.75 were set and a false discovery rate of 0.01 was used. Quantification was done using the protein LFQ method Quant 2.0 (SN standard) within Spectronaut.

Output proteomics files were handled in R Studio (version 2026.01.0). Raw proteomics intensities were median-normalised across samples followed by filtering to exclude low-information proteins (rows containing only 0/1 peptide counts). Data were log2-transformed and further filtered to retain proteins with sufficient observations (minimum of 70% of the values should be numerical least one experimental group), with remaining missing values imputed using group-wise row means and duplicate gene entries collapsed by averaging. Differential expression between disease and control or WT and TG groups at each time point was assessed using Welch’s t-tests, with log2 fold change calculation and multiple testing correction performed using the Benjamini–Hochberg false discovery rate.

Volcano plot of gene expression data was generated using ggplot2 package, applying thresholds of minimum of 20% biological change: log2 foldchange > 0.3 or < −0.3 and −log10(p-value) > 1.3 (≈ p < 0.05) to classify genes as up- or downregulated also called differentially expressed (DE) proteins. Significant genes were colour-coded and labelled using ggrepel package. Heatmaps were plotted using pheatmap package.

#### Weighted Gene Co-expression Network Analysis (WGCNA)

WGCNA analysis was performed using WGCNA R package (Langfelder & Horvath, 2008). Prior to network construction samples were quality checked and sample clustering was performed to detect outliers (Suppl fig X.). To construct a scale-free network, an appropriate soft-thresholding power (β) was selected using the pickSoftThreshold function. The scale-free topology fit index (R²) was calculated across a range of powers. A power of β = 16 was selected for human data and β = 11 for the hTDP overexpressed rodent model to ensure ideal scale-free topology while maintaining sufficient mean connectivity (Supplfig X). The resulting signed network was used for all downstream analyses. A weighted gene co-expression network was constructed by calculating pairwise correlations between all genes using bicor function, which were then raised to the soft-thresholding power to generate an adjacency matrix. The adjacency matrix was transformed into a Topological Overlap Matrix (TOM) to capture shared network connections between gene pairs, and the corresponding dissimilarity matrix (1 − TOM) was used for hierarchical clustering. Gene modules were identified using the cutreeDynamic function with the minimum module size of 20 genes. Genes that could not be assigned to any module were placed in the grey module and excluded from further analyses.

#### Module–Trait Correlation Analysis

To identify modules associated with clinical traits, such as genetics or disease progression in case of the rodent model, biweight midcorrelation (bicor) correlations were calculated between each module eigengene and each trait. A correlation heatmap was generated displaying both the correlation coefficient (r) and associated p-value for each module–trait pair. Modules with p < 0.05 were considered significantly associated with a given trait and selected for further analysis. For each gene, two key metrics were calculated to quantify its biological relevance: Gene Significance (GS) which is the absolute Pearson correlation between each gene’s expression profile and the trait of interest. Higher GS indcates a stronger direct association with the trait. Module Membership (MM / kME) was measured using Pearson correlation between each gene’s expression profile and the module eigengene. Higher MM indicates a gene is more central to its module (i.e. a hub gene). GS was calculated for each trait independently, yielding a GS matrix across all genes and traits. MM was calculated using the signedKME function. Hub genes were defined as genes with both high Module Membership and high Gene Significance for the trait associated with their module. For each significant module, genes were ranked by a combined score calculated as the average of absolute MM and GS visualising genes per module were retained as hub genes and used for generating Chord diagram using the TOM-based connection weight.

#### Gene enrichment analysis (GSEA)

To characterise the biological functions of significant modules, gene ontology (GO) and pathway enrichment analyses (REACTOME, KEGG – REFS) were performed using the clusterProfiler package. Enrichment was tested against a custom background (all identified proteins in the corresponding dataset) using Benjamini–Hochberg FDR correction. Enriched themes were visualised using the ggplot2 package.

#### 6-plex TMT proteomics

Raw MaxQuant intensity profiles were filtered to retain proteins identified with ≥2 unique peptides and detected across all samples, yielding 5,244 analysis-ready proteins. ALS data were normalised to controls and further median-normalised to correct for minor loading differences, generating ratiometric protein expression values. Values were log2 transformed and Z-scores were counted as shown before (Laszlo 2022). A threshold of ±20% change (≥1.2 or ≤0.8) was applied to define biologically relevant differences, consistent with the sensitivity of downstream validation methods.

To identify shared and brain area and disease-specific functional profiles, differentially expressed proteins were stratified into distinct directional groups based on differentially expressed genes (log2FC values) per BA17, BA4 and BA9 or C9neg, C9pos, APoE3 and APoE4 genotypes. Functional enrichment analysis was performed using the gprofiler2 package in R, applying an adjusted significance cutoff of (p < 0.05). Finally, the top five enriched biological terms from each profile were integrated and visualised using a comprehensive, shared-axis dot plot to trace convergent and divergent pathological mechanisms across both disorders.

### Array tomography

Array tomography was performed as previously described^2,3^. Briefly, fresh post-mortem tissue from BA17/19 was fixed in 4% paraformaldehyde in PBS for 2–3 h, dehydrated through graded ethanol, and embedded in LR White resin prior to polymerisation at 60 °C. Ultrathin serial sections (70 nm) were cut using an ultramicrotome and collected as ribbons on gelatin-coated coverslips. Sections were permeabilised (50 mM glycine) and blocked (0.1% fish skin gelatin, 0.05% Tween-20 in TBS) prior to overnight incubation with primary antibodies against synaptophysin (1:50). Fluorescent secondary antibodies (Alexa Fluor 488, 1:50) and DAPI were applied the following day, and samples were mounted for imaging. Image stacks were acquired at consistent positions along each ribbon using a DeltaVision Elite widefield fluorescence microscope with a 63×/1.4 NA objective. At least two stacks per region per case were collected. Stacks were aligned using the ImageJ MultiStackReg plugin, followed by thresholding, segmentation, and puncta density quantification using a custom MATLAB pipeline (Extended Data Figure 2). Analysis tools publicly available via GitHub (https://github.com/arraytomographyusers/Array_tomography_analysis_tool).

### Electron microscopy

Transmission electron microscopy was performed as previously described^2,41^. Briefly, post-mortem cortical samples were fixed in 4% paraformaldehyde and 2.5% glutaraldehyde in 0.1 M PB, processed with osmium tetroxide, dehydrated, and embedded in Durcupan resin. Ultrathin sections (70 nm) were collected on grids, stained with lead citrate, and imaged using a Philips CM12 transmission electron microscope equipped with a Gatan digital camera. Synaptic profiles were analysed by an investigator blinded to disease status and identified by the presence of defined pre- and post-synaptic compartments containing synaptic vesicles and a postsynaptic density, respectively. Degenerating synapses were classified based on electron-dense cytoplasmic morphology as previously described^42^. Axonal profiles were analysed by electron microscopy in five distinct white matter regions across the brain (Figure 1g). Healthy axons were identified based on the presence of compact myelin sheaths and normal axoplasmic morphology. The number of healthy axonal profiles (myelin rings) was counted within each region of interest, and axonal density was calculated by normalising the counts to the analysed area (µm²).

### Statistical analysis

Group-wise comparisons were tested for statistical significance using Prism 10 (GraphPad). Normality was measured using Shapiro-Wilk normality test or Kolmogorov-Smirnov test. Unpaired comparisons were analyzed using two-sided Student’s t tests (normally distributed) and by Mann–Whitney U tests (not normally distributed). The level of significance was set to P < 0.05 in every case; exact P values and the type of test performed are indicated in each figure legend. Micrographs in the figures are shown as representative images. No statistical methods were used to predetermine the sample size; it was defined by the availability of human samples.

**Supplementary Table 3.**
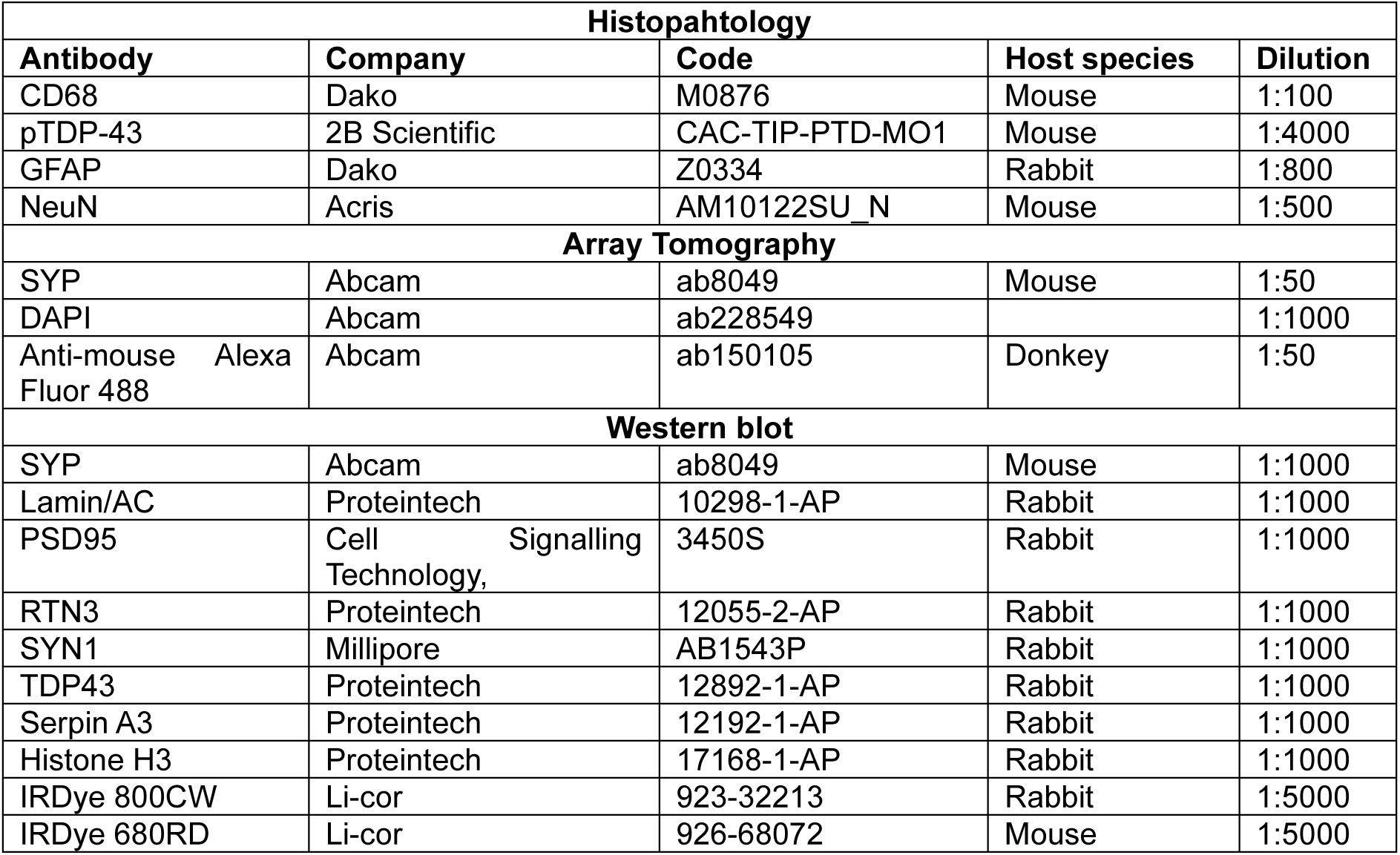
Primary and secondary antibody information.

## Acknowledgements

The authors wish to thank the donors and their families for the generous tissue donations and the Edinburgh Brain Bank team for processing and providing materials, especially to facility manager Karina McDade and director, Prof. Colin Smith. The authors acknowledge the Dundee Proteomics FingerPrints Facility, namely, Douglas Lamont, Samantha Kosto and Alan Score, for their technical support in processing samples for MS experiments. The authors wish to thank the staff of the Dundee Imaging Facility and the Glasgow Tissue Research facility. We acknowledge input of Dr Wei Luan for animal tissue collection and data interpretation. The research was funded by The Neurosciences Foundation and the Motor Neurone Disease Association by supporting Z.I.L. with a Junior Non-Clinical Fellowship (Oct21/977-799). Research was also funded by the Alzheimer’s Society by supporting C.M.H. with a Dementia Research Leader Fellowship, and the Academy of Medical Sciences funding C.M.H. with a Springboard Award. We are grateful for the Euan MacDonald Centre for Motor Neuron Disease Research funding A.S.A.’s PhD studies. The study was also funded by FightMND funding A.K.W. with a Bill Guest Mid-Career Research Fellowship, the Ross Maclean Fellowship and Brazil Family Program for Neurology funding to A.K.W., and the UK Dementia Research Institute (grant UKDRI-Edin005, UKDRI-4004 to T.L.S.-J.), through UK DRI, principally funded by the UK Medical Research Council.

## Author Contributions

C.M.H. conceived and oversaw the study. Z.I.L. designed, performed and analysed the proteomics studies. Z.I.L. analysed histological samples. Array tomography was performed by A.S.A. and analysed by Z.I.L. T.H.G. collected the TEM images, which were analysed by Z.I.L. and A.S.A. Western blot experiments were performed by A.M. and Z.I.L. The hTDP-43/rNLS8 rodent tissue was collected and provided by R.S.G and A.K.W. Mouse synaptoneurosomes were generated by D.vd.H. and Z.I.L. TLSJ provided Alzheimer’s disease proteomics data. Z.I.L. and C.M.H. wrote the paper. All authors contributed to editing and reviewing the paper and approve of its publication.

## Competition of interests

TSJ has received compensation for consulting, grant reviews, scientific talks, or collaborative research over the past 10 years from AbbVie, Sanofi, Merck, Autifony, Scottish Brain Sciences, Jay Therapeutics, Cognition Therapeutics, Ono, Novo Nordisk, Eisai, Boehringer Ingelheim, and Bristol-Myers Squibb. THG has provided advisory services to LifeArc, Novartis and Roche.

## Data Availability

The mass spectrometry proteomics data generated in this study have been deposited in the PRIDE repository (ProteomeXchange Consortium) under accession number [*provided upon publication*]. All other data supporting the findings of this study are available from the corresponding author upon reasonable request.

## Extended Data Figures

**Extended Data Figure 1.**
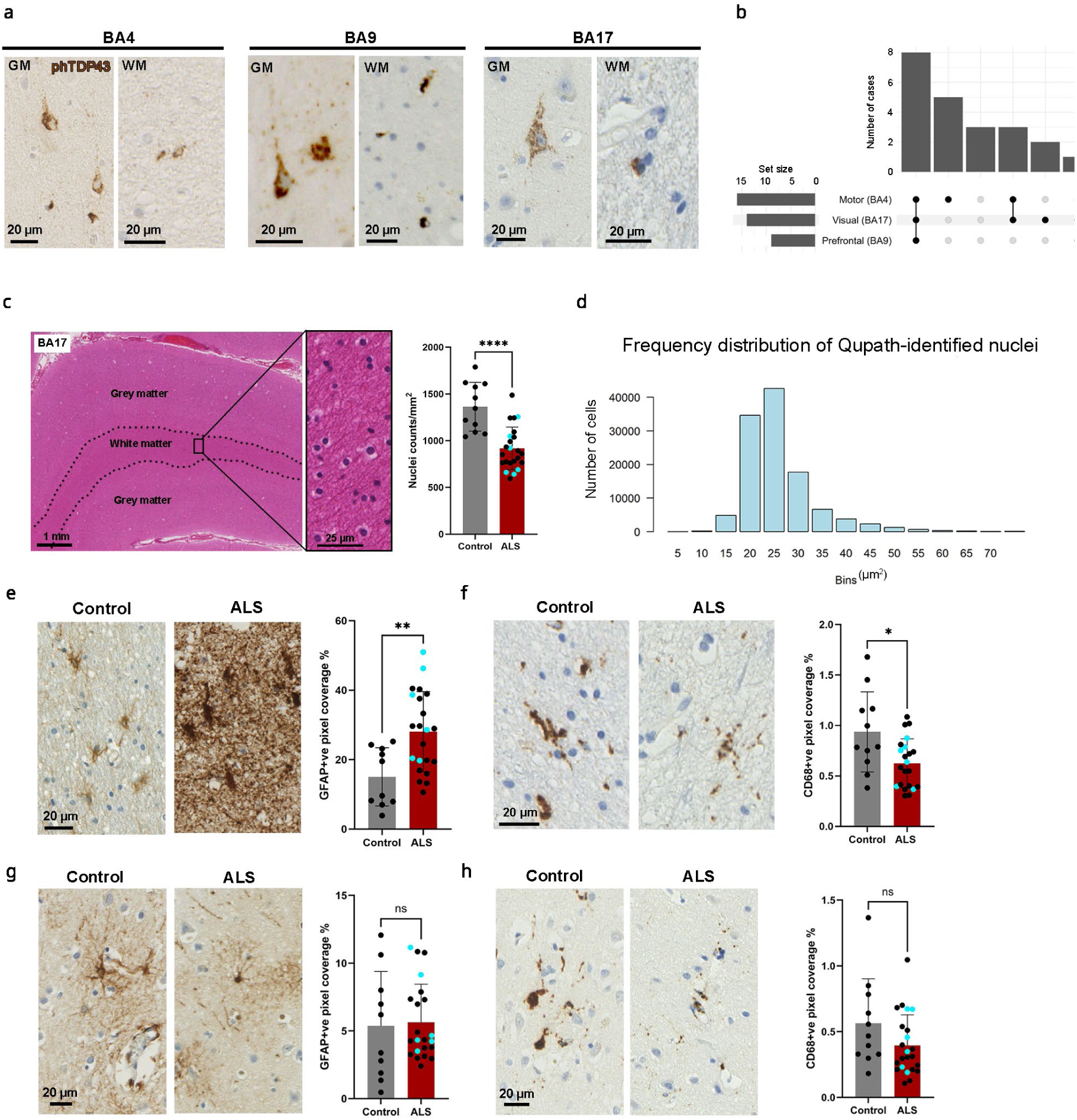
QuPath-based digital pathology analysis of cellular changes in BA17. **a**, Representative pTDP43 immunohistochemical staining demonstrating aggregated pathology across BA4, BA9, and BA17 cortical areas within the same donor (GM=grey and WM=white matter). **b**, UpSet plot showing the intersection of pTDP43 pathology distribution across the analyzed brain regions (n=23). **c**, Example cortical H&E-stained section and quantification of cell density within the BA17-associated ALS white matter. Welch’s t test p<0.0001, Ctrl n=11, ALS n=23. **d**, Frequency distribution of cell nuclei diameters identified via QuPath automated Watershed cell detection. **e–h**, Histological quantification of GFAP-positive astrocyte (**e,g)** and CD68-positive microglia (**f,h**) pixel coverage in BA17/19 white matter (**e,f)** and grey matter (**g,h**). Welch’s t test (**e**) p=0.0015, Ctrl n=10, ALS n=22, (**f**) p=0.0318, Ctrl n=11, ALS n=22, (**g**) Mann Whitney test, p=0.5087, Ctrl n=10, ALS n=22, (**h**) p=0.1233, Ctrl n=11, ALS n=23. Cyan symbols indicate *C9orf72*-positive cases. Bar graphs show mean±SD

**Extended Data Figure 2.**
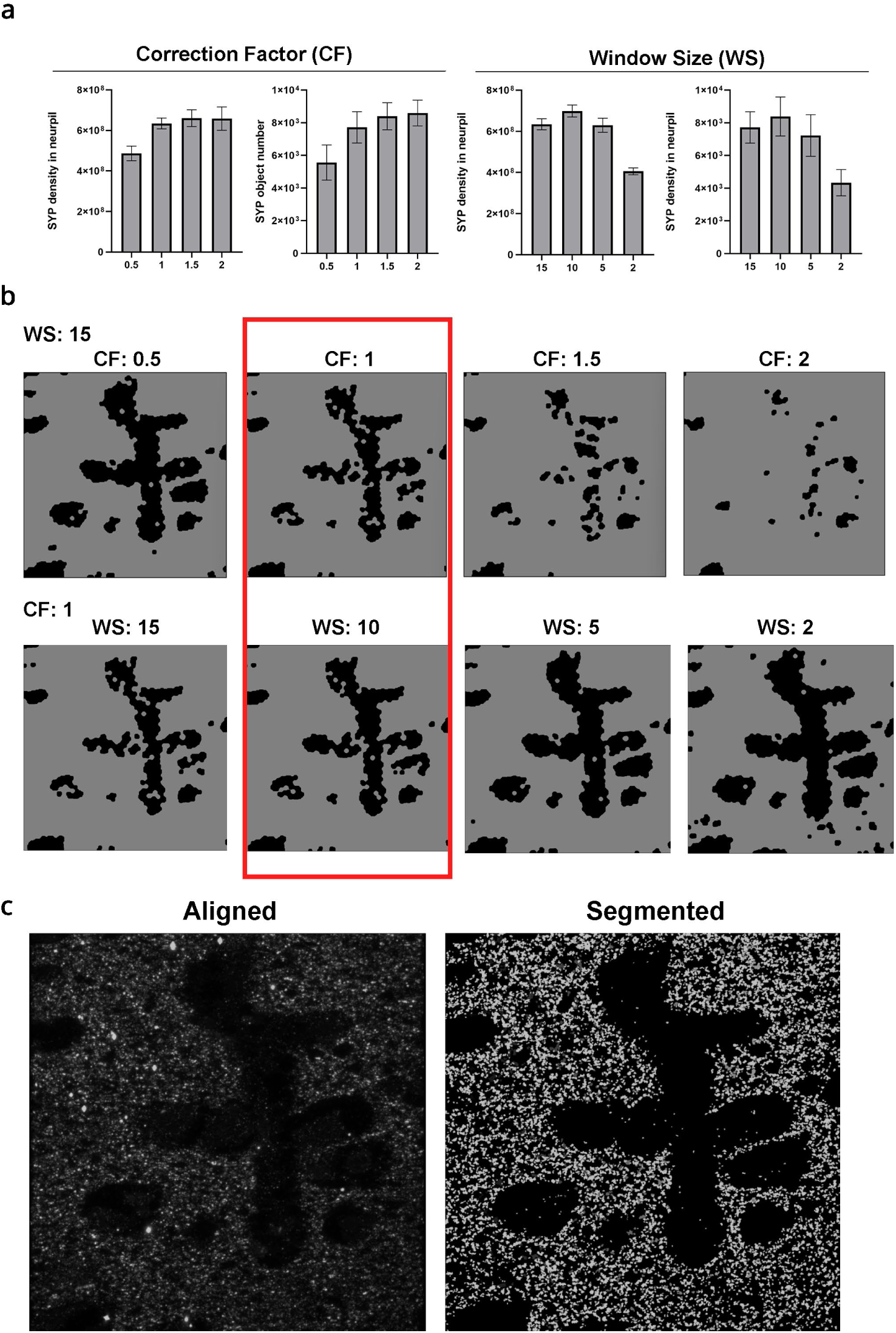
Image segmentation optimization parameters for array tomography. **a**, Evaluation of SYP density quantification across varying correction factors (CF) and window sizes (WS). **b**, Representative segmentation masks generated using different CF values at a fixed WS, and varying WS values at a fixed CF (CF = 1). **c**, Aligned and segmented example 3D-images show synaptic puncta isolation for further analysis.

**Extended Data Figure 3.**
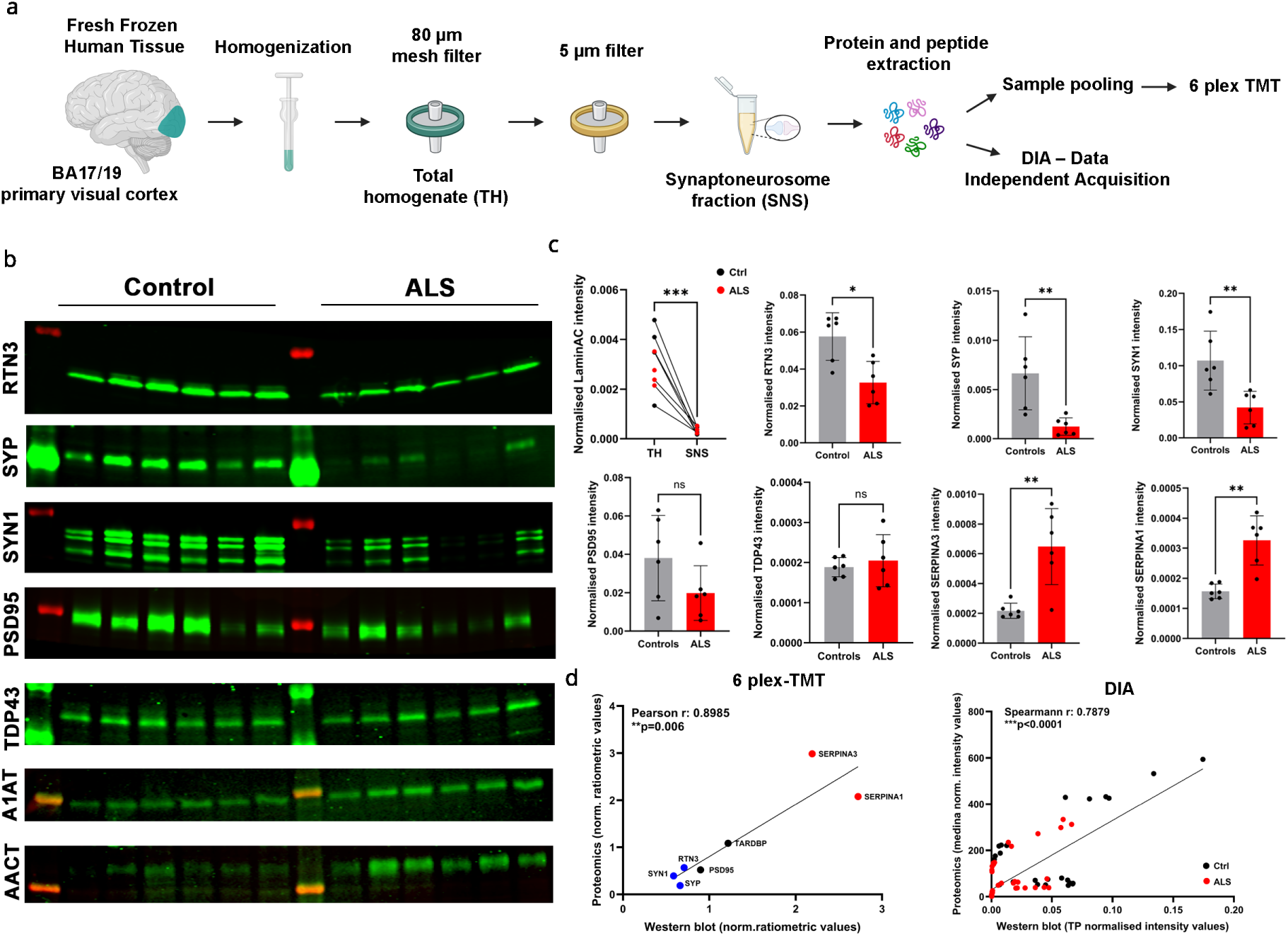
Proteomic validation and quality control of isolated synaptoneurosomes. **a**, Methodological schematic of the synaptoneurosome (SNS) preparation and downstream liquid chromatography–mass spectrometry (LC-MS/MS) proteomic workflows. **b**, Representative western blot images for validation targets selected across proteomic expression ranges (LaminAC: tailed-paired t test p=0.0002, number of pairs = 8. RTN3 Mann Whitney test p= 0.0152; PSD95: Unpaired t test p= 0.1218; SYP: Mann Whitney test p=0.0043; TDP-43: Mann Whitney test p>0.9999; SYN1: Unpaired t test p=0.0066; SerpinA3: Mann Whitney test 0.0087; SerpinA1 Mann Whitney test p=0.0022; Ctrl n=6 and ALS n=6). **c**, Densitometric quantification of western blot band intensities. **d**, Correlation analysis between western blot normalized intensities and median-normalized proteomic abundances from 6-plex TMT and DIA datasets.

**Extended Data Figure 4.**
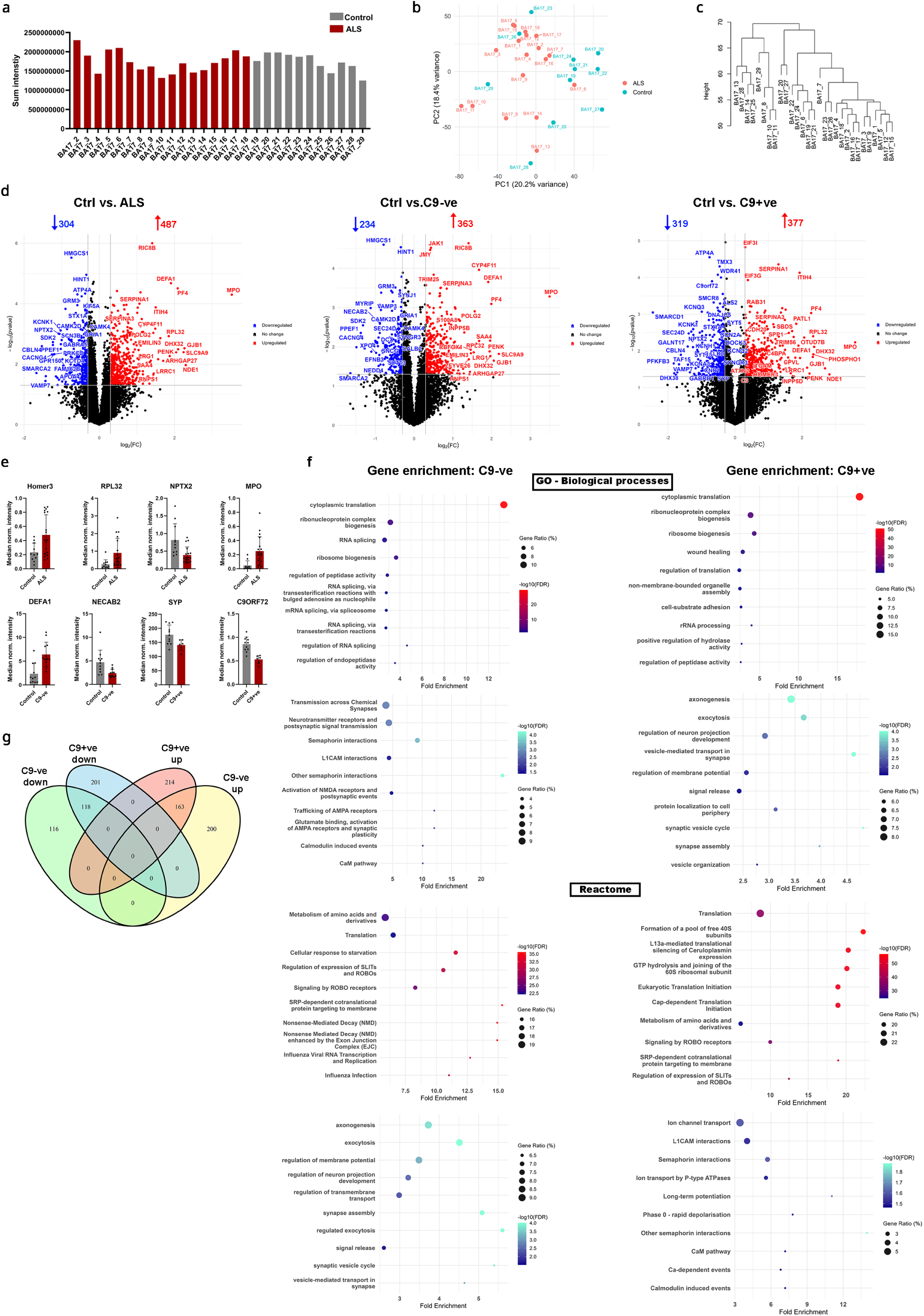
Bioinformatic quality control and differential expression analysis of visual cortex DIA proteomics. **a**, Summed MS2 intensity distributions per sample demonstrating overall signal consistency. **b**, Principal component analysis (PCA) score plot showing sample clustering based on global proteomic profiles. **c**, Unsupervised hierarchical clustering dendrogram of all analyzed samples. **d**, Volcano plots of differentially expressed proteins (DEPs) across the indicated experimental groups. Thresholds: –log₁₀(p) ≥ 1.35 (p < 0.05) and |log₂FC| ≥ 0.3 (≥20% biological change); arrows indicate DEP counts. **e**, Sample-level variance analysis across comparisons validating the statistical framework. **f**, Gene Ontology (GO) enrichment analysis of up- and downregulated proteins in control vs. *C9orf72*-negative and control vs. *C9orf72*-positive comparisons. **g**, Venn diagram showing the overlap of DE proteins across genotype groups.

**Extended Data Figure 5.**
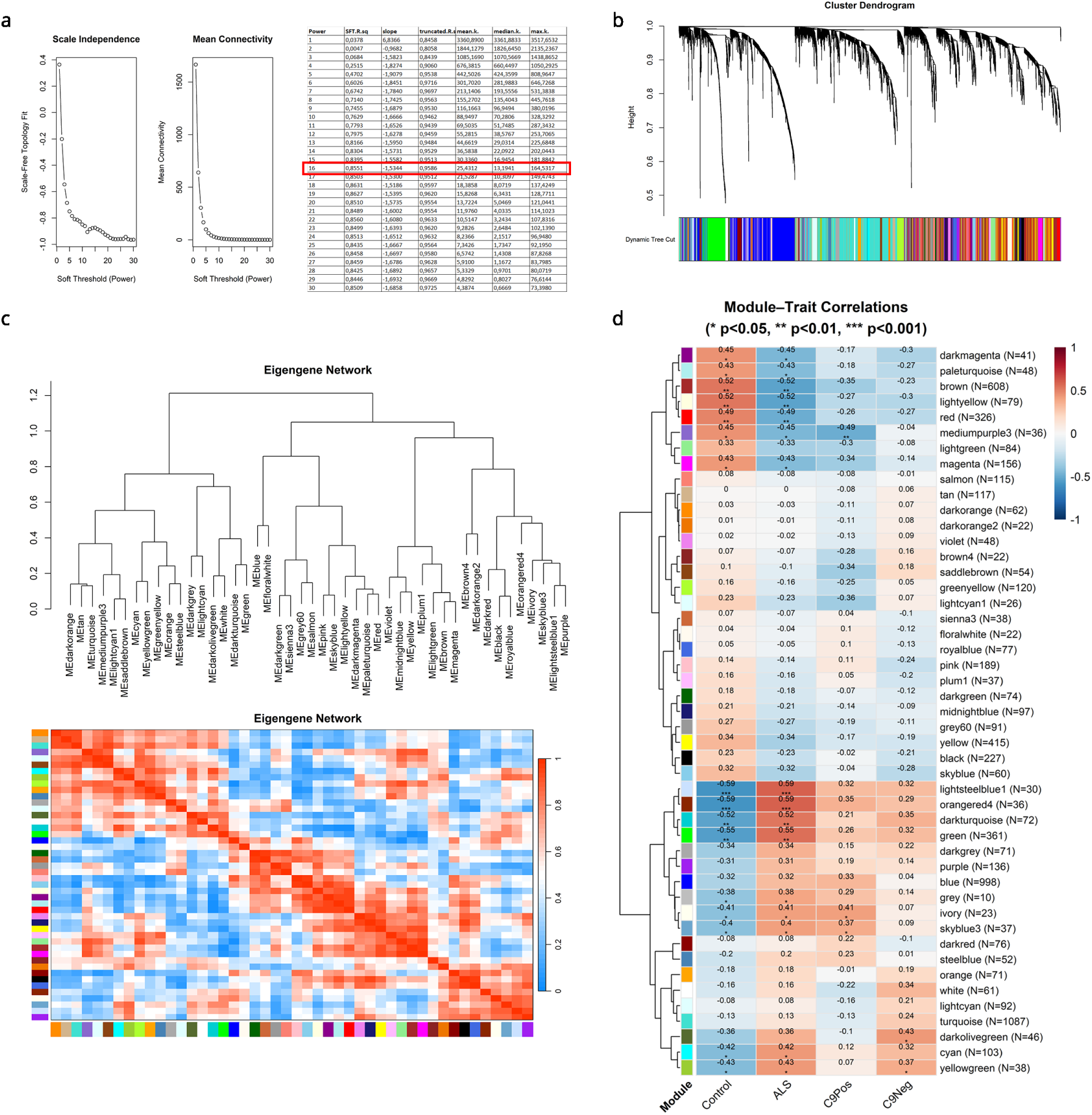
WGCNA network construction parameters and module structure. **a**, Soft-thresholding power selection based on scale-free topology (SFT) criteria. SFT index (R²) and mean connectivity were evaluated across powers (1–30) using biweight midcorrelation; the power threshold was selected where R² ≥ 0.8–0.9. **b**, Hierarchical clustering dendrogram of proteins with dynamic tree cut module assignments indicated by color bars. **c**, Module eigengene clustering and adjacency heatmap displaying inter-module relationships. **d**, Complete module–trait correlation heatmap with significance values: · p < 0.1, * p < 0.05, ** p < 0.01, *** p < 0.001.

**Extended Data Figure 6.**
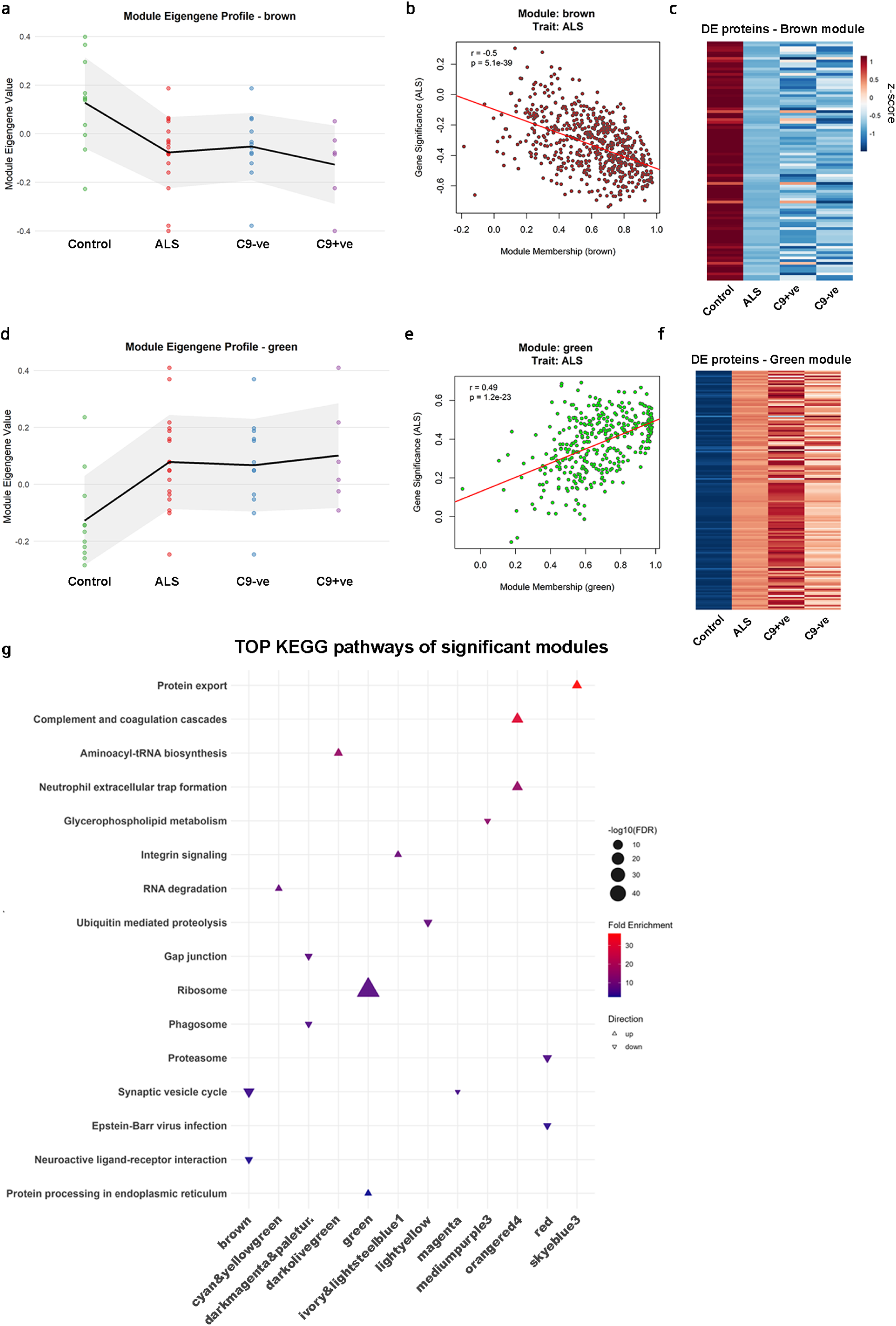
Characterization of top trait-associated WGCNA modules. **a,d**, Spaghetti plots showing module eigengene values per sample across disease groups for the brown (**a**) and green (**d**) modules. Black lines indicate group means ± s.d. (grey shading). **b,e**, Scatter plots of gene significance (GS) versus module membership (MM) for the brown (**b**) and green (**e**) modules. **c,f**, Expression heatmaps of individual DEPs within the selected modules across all samples. **g**, Gene Set Enrichment Analysis (GSEA) normalized enrichment scores for the top functional terms.

**Extended Data Figure 7.**
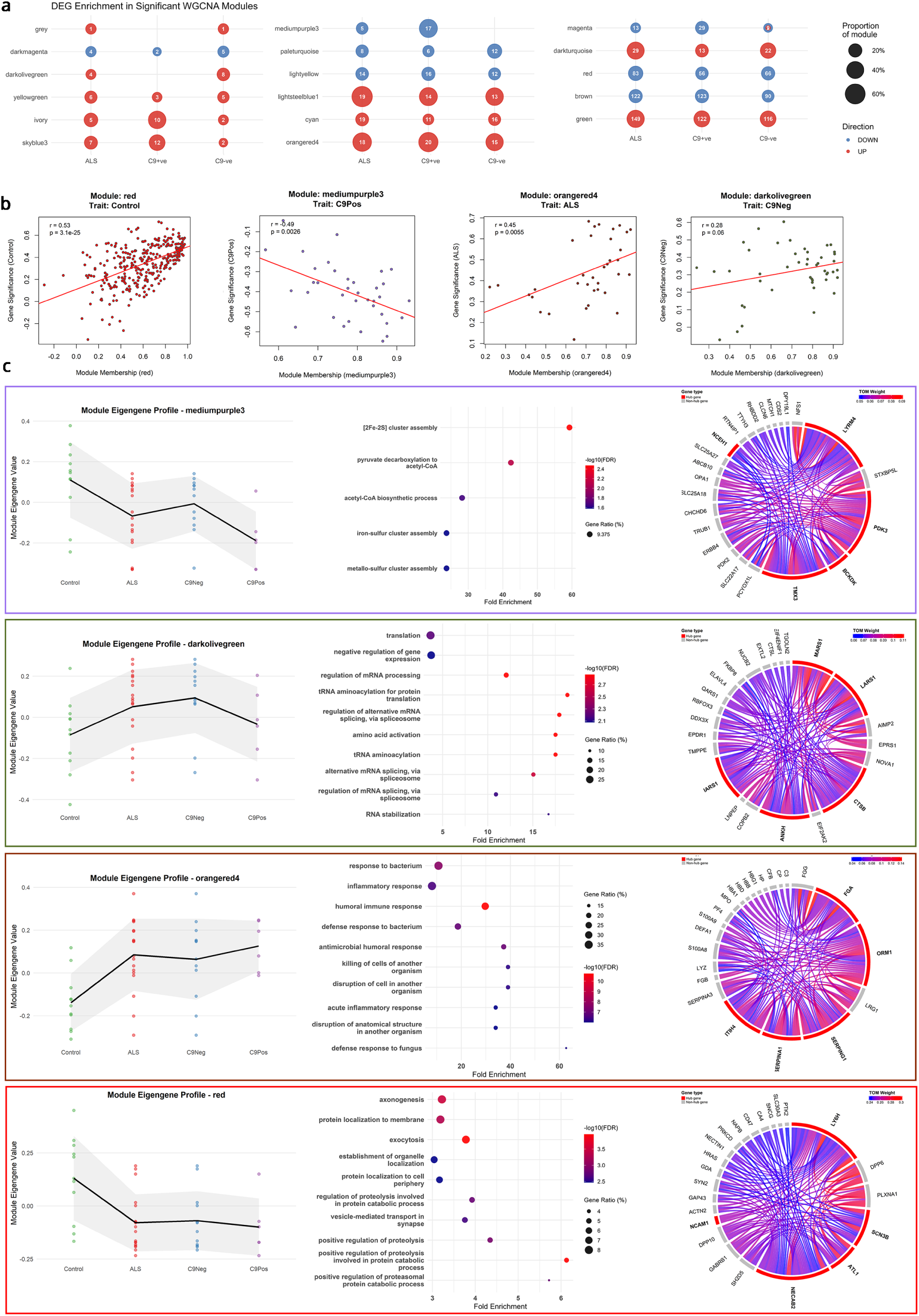
ALS molecular subtype-specific WGCNA network analysis. **a**, Bubble plot displaying the number of DEPs mapping to individual WGCNA modules. **b**, Correlation plots between GS and MM across disease-subtype networks. **c**, Module eigengene profiles, GO Biological Process, GSEA enrichment terms, and chord diagrams displaying top 5 hub protein topologies for the selected subtype-specific modules.

**Extended Data Figure 8.**
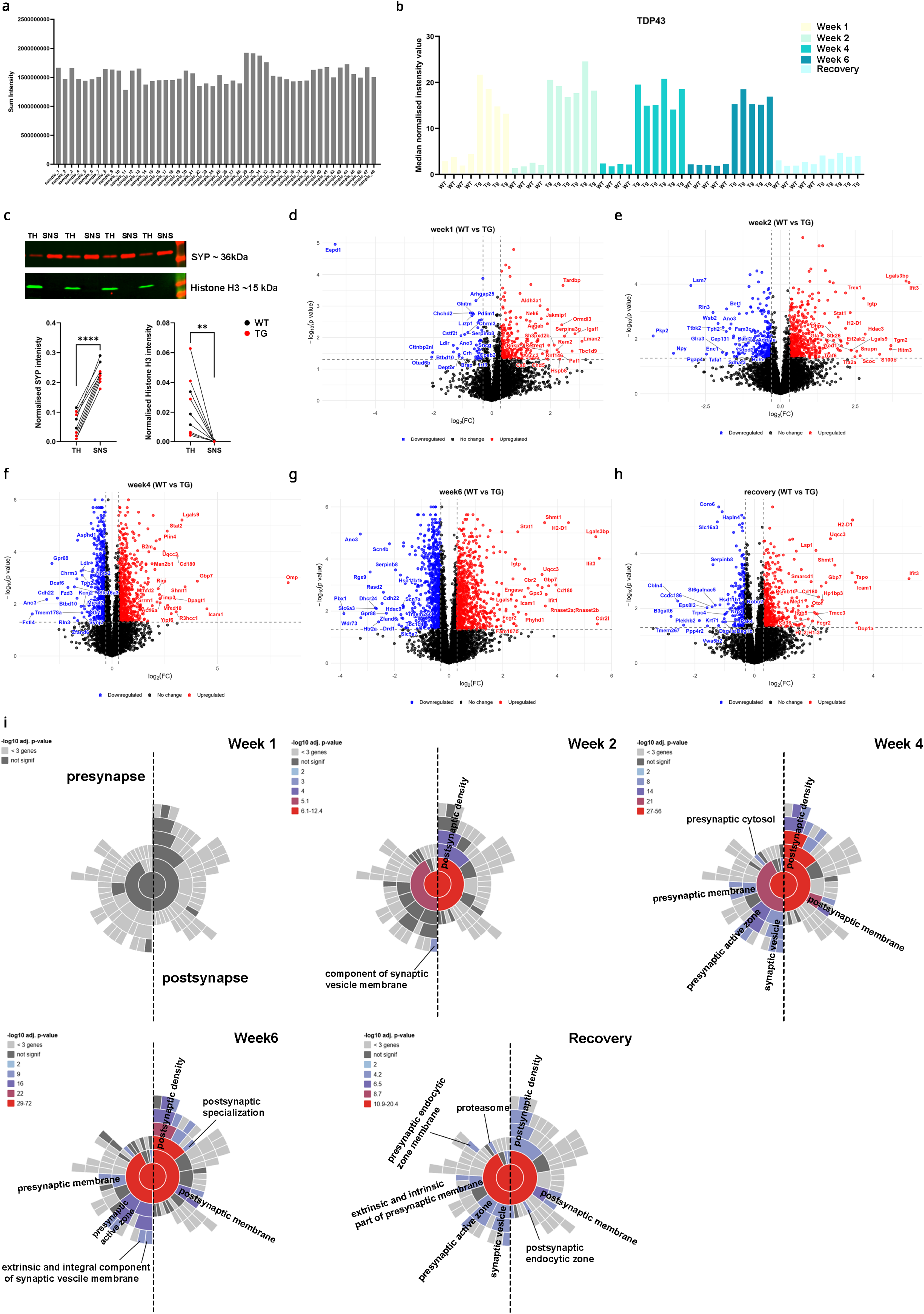
Differential protein expression profiles across longitudinal timepoints in the hTDP-43 rNLS8 mouse model. **a**, Raw summed MS2 intensity profiles per sample. **b**, Normalized TDP-43 intensity values confirming transgenic overexpression in TG animals. **c**, Representative western blot images and densitometric quantification confirming synaptophysin (SYP) enrichment in SNS fractions and exclusion of the nuclear marker histone H3 (HH3) (two-tailed paired t-test, SYP p<0.0001, HH3 p=0.007, number of pairs = 9). **d–h**, Volcano plots from multiple t-tests with Benjamini–Hochberg correction identifying DE proteins across the indicated timepoint comparisons. Red points indicate upregulated proteins; blue points indicate downregulated proteins (thresholds: –log₁₀(p) ≥ 1.35; |log₂FC| ≥ 0.3)). **i**, Sunburst plots illustrating the progressive enrichment of pre- and postsynaptic functional terms from week 1 through week 6, with persistent compartment alterations during the recovery stage.

**Extended Data Figure 9.**
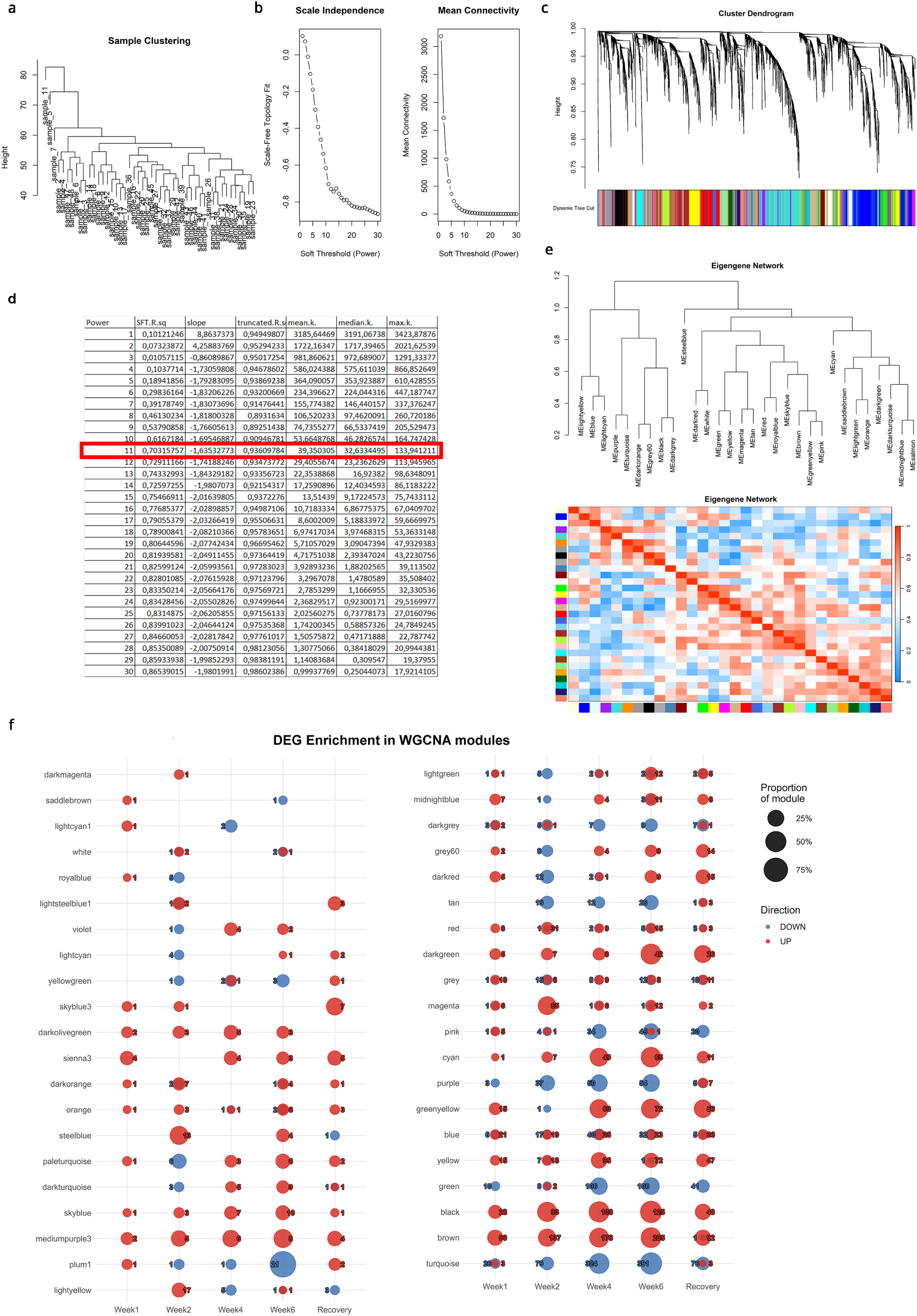
WGCNA network parameters and module architecture for the hTDP-43 rNLS8 mouse model. **a**, Unsupervised hierarchical clustering of individual mouse proteomic samples. **b,d**, Soft-thresholding power selection based on scale-free topology (SFT) index (R² ≥ 0.8) (**b**) and mean connectivity (**d**) evaluated across powers 1–30 using biweight midcorrelation. **c**, Hierarchical clustering dendrogram of proteins with dynamic tree cut module assignments indicated by colour bars. **e**, Module eigengene clustering and adjacency heatmap displaying inter-module eigengene correlations. **f**, Total number of downregulated (blue) and upregulated (red) DE proteins mapping to individual WGCNA modules across timepoints, identified via multiple t-tests.

**Extended Data Figure 10.**
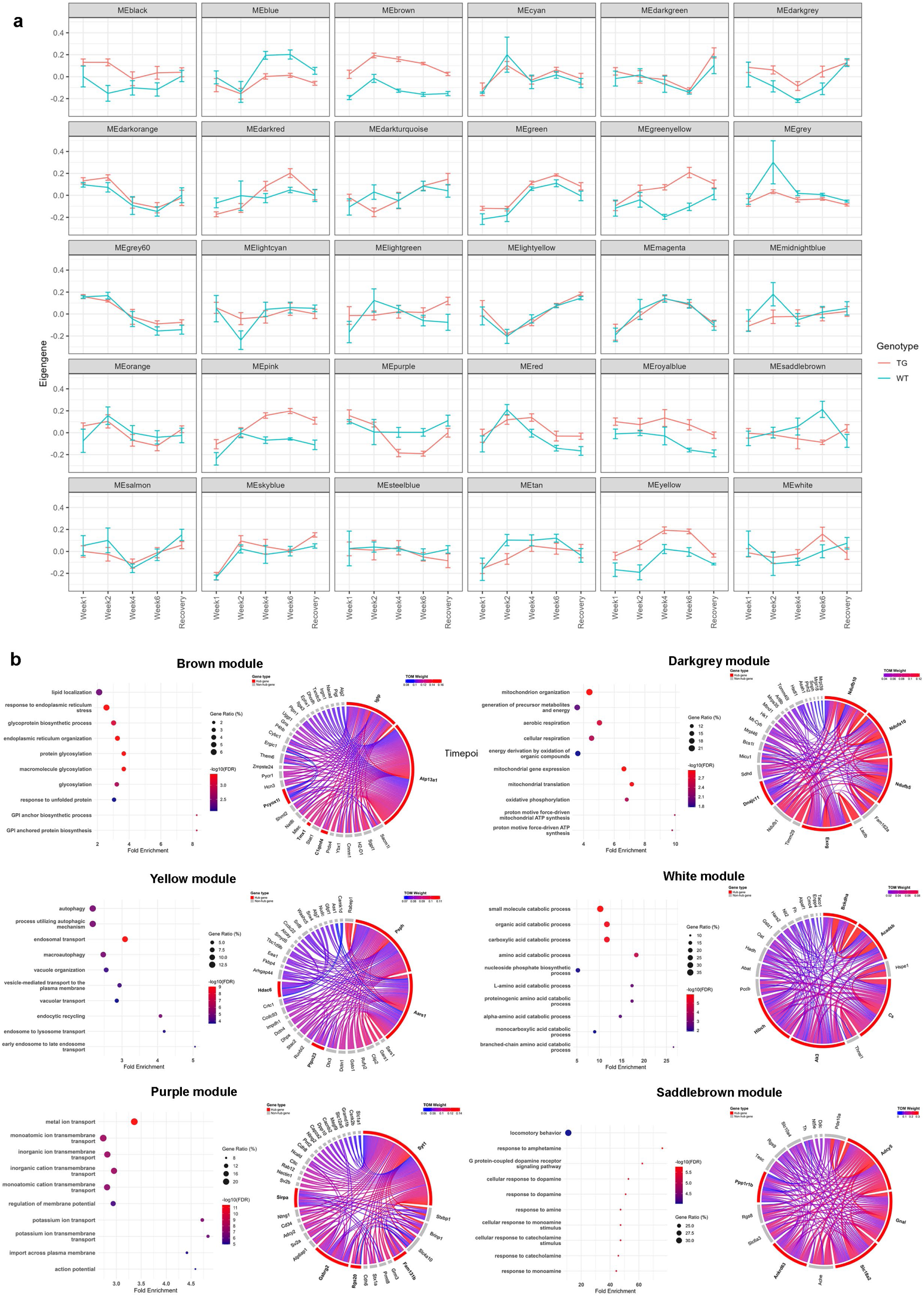
Additional WGCNA module trajectories and enrichment profiles from the hTDP-43 rNLS8 mouse model. **a**, Spaghetti plots showing genotype-stratified sample trajectories across additional co-expression modules. **b**, Characterization of selected WGCNA modules showing the top enriched GO Biological Process terms from GSEA, alongside chord diagrams illustrating the network topology of the top 5 hub genes and their closest co-expression connections.

**Extended Data Figure 11.**
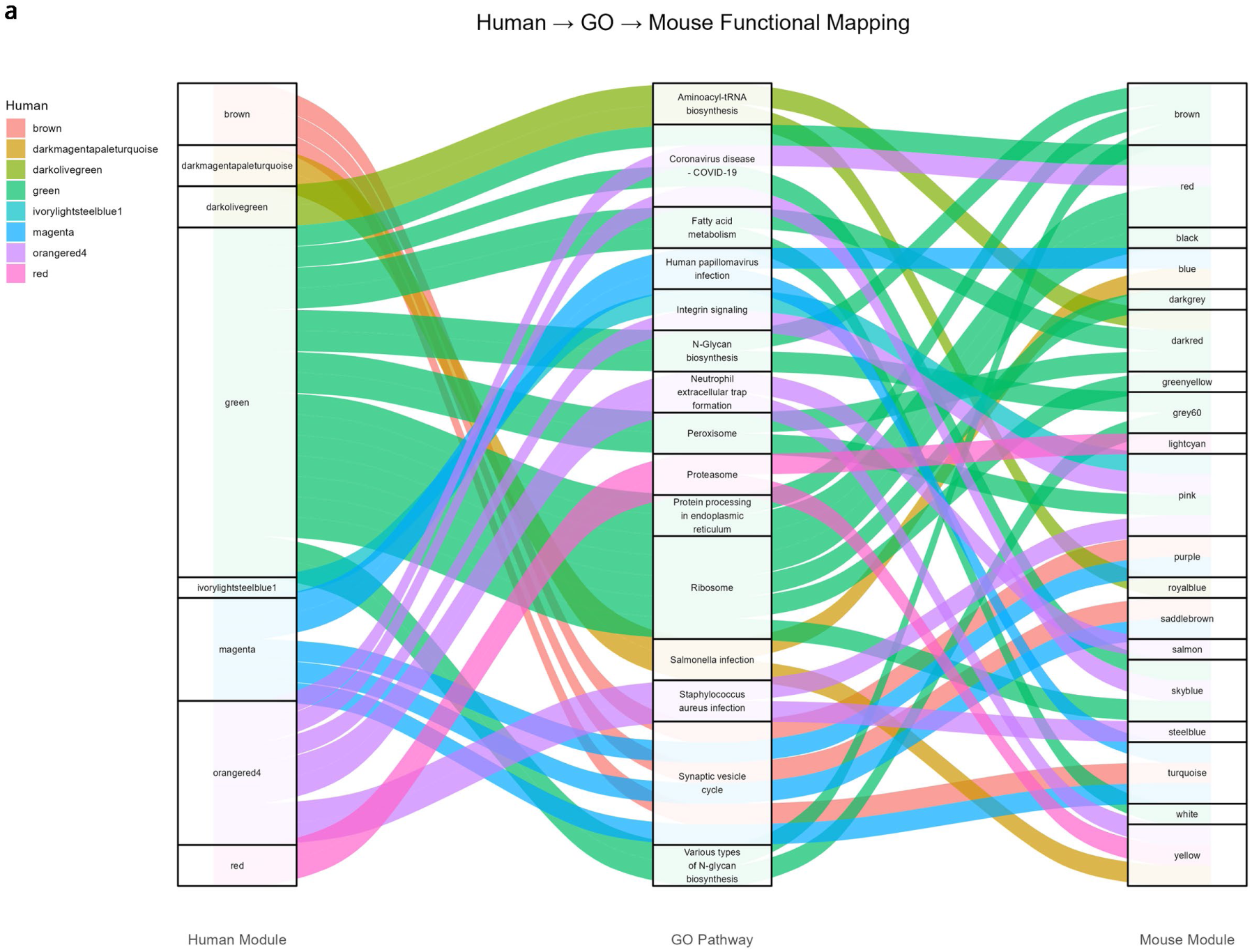
Conservation of functional protein enrichment themes across species. Alluvial plot illustrating the shared GO Biological Process functional themes and mapping alignments between human and mouse WGCNA co-expression modules.

## Notes

https://henstridgelab.shinyapps.io/mnd-synaptic-proteomics/

## References

1. Fogarty, M. J., Noakes, P. G. & Bellingham, M. C. Motor cortex layer V pyramidal neurons exhibit dendritic regression, spine loss, and increased synaptic excitation in the presymptomatic hSOD1G93A mouse model of amyotrophic lateral sclerosis. Journal of Neuroscience 35, 643–647 (2015).

2. Henstridge, C. M. et al. Synapse loss in the prefrontal cortex is associated with cognitive decline in amyotrophic lateral sclerosis. Acta Neuropathol. 135, 213–226 (2018).

3. Laszlo, Z. I. et al. Synaptic proteomics reveal distinct molecular signatures of cognitive change and C9ORF72 repeat expansion in the human ALS cortex. Acta Neuropathol. Commun. 10, 1–20 (2022).

4. Schanz, O. et al. Cortical hyperexcitability in patients with C9ORF72 mutations: Relationship to phenotype. Muscle Nerve 54, 264–269 (2016).

5. Aly, A. et al. Integrative proteomics highlight presynaptic alterations and c-Jun misactivation as convergent pathomechanisms in ALS. Acta Neuropathol. 146, 451–475 (2023).

6. Silva-Hucha, S. et al. Excitotoxicity in amyotrophic lateral sclerosis: a key pathogenic mechanism. Brain Communications vol. 8 Preprint at 10.1093/braincomms/fcag098 (2026).

7. Sommer, D. et al. Aging-Dependent Altered Transcriptional Programs Underlie Activity Impairments in Human C9orf72-Mutant Motor Neurons. Front. Mol. Neurosci. 15, (2022).

8. Clayton, E. L., Huggon, L., Cousin, M. A. & Mizielinska, S. Synaptopathy: presynaptic convergence in frontotemporal dementia and amyotrophic lateral sclerosis. Brain vol. 147 2289–2307 Preprint at 10.1093/brain/awae074 (2024).

9. Perkins, E. M. et al. Altered network properties in C9ORF72 repeat expansion cortical neurons are due to synaptic dysfunction. Mol. Neurodegener. 16, (2021).

10. Scekic-Zahirovic, J., et al. Cortical Hyperexcitability in Mouse Models and Patients with Amyotrophic Lateral Sclerosis Is Linked to Noradrenaline Deficiency. Sci. Transl. Med vol. 16 https://www.science.org (2024).

11. Haidar, M. et al. Cortical hyperexcitability drives dying forward amyotrophic lateral sclerosis symptoms and pathology in mice. Prog. Neurobiol. 252, (2025).

12. Chipika, R. H. et al. Alterations in somatosensory, visual and auditory pathways in amyotrophic lateral sclerosis: an under-recognised facet of ALS. J. Integr. Neurosci. 21, (2022).

13. Zou, T. et al. Medulla oblongata dominated synaptic density network degeneration in amyotrophic lateral sclerosis. Neuroimage Clin. 47, (2025).

14. Brettschneider, J. et al. Stages of pTDP-43 pathology in amyotrophic lateral sclerosis. Ann. Neurol. 74, 20–38 (2013).

15. Braak, H. et al. Amyotrophic lateral sclerosis - A model of corticofugal axonal spread. Nature Reviews Neurology vol. 9 708–714 Preprint at 10.1038/nrneurol.2013.221 (2013).

16. Cykowski, M. D., Arumanayagam, A. S., Powell, S. Z. & Appel, S. H. Primary visual cortex pathology in ALS patients with C9ORF72 expansion. Brain Pathology vol. 34 Preprint at 10.1111/bpa.13229 (2024).

17. Porta, S. et al. Distinct brain-derived TDP-43 strains from FTLD-TDP subtypes induce diverse morphological TDP-43 aggregates and spreading patterns in vitro and in vivo. Neuropathol. Appl. Neurobiol. 47, 1033–1049 (2021).

18. Porta, S. et al. Patient-derived frontotemporal lobar degeneration brain extracts induce formation and spreading of TDP-43 pathology in vivo. Nature Communications 9, (2018).

19. Walker, A. K. et al. Functional recovery in new mouse models of ALS/FTLD after clearance of pathological cytoplasmic TDP-43. Acta Neuropathol. 130, 643–660 (2015).

20. Spiller, K. J. et al. Selective motor neuron resistance and recovery in a new inducible mouse model of TDP-43 proteinopathy. Journal of Neuroscience 36, 7707–7717 (2016).

21. Luan, W. et al. Early activation of cellular stress and death pathways caused by cytoplasmic TDP-43 in the rNLS8 mouse model of ALS and FTD. Mol. Psychiatry 28, 2445–2461 (2023).

22. San Gil, R., et al. A transient protein folding response targets aggregation in the early phase of TDP-43-mediated neurodegeneration. Nat. Commun. 15, (2024).

23. Hesse, R. et al. Comparative profiling of the synaptic proteome from Alzheimer’s disease patients with focus on the APOE genotype. Acta Neuropathol. Commun. 7, 1–18 (2019).

24. Eisen, A., Vucic, S. & Mitsumoto, H. History of ALS and the competing theories on pathogenesis: IFCN handbook chapter. Clinical Neurophysiology Practice vol. 9 1–12 Preprint at 10.1016/j.cnp.2023.11.004 (2024).

25. Eisen, A. The dying forward hypothesis of als: Tracing its history. Brain Sciences vol. 11 1–9 Preprint at 10.3390/brainsci11030300 (2021).

26. Maranzano, A. et al. Regional spreading pattern is associated with clinical phenotype in amyotrophic lateral sclerosis. Brain 146, 4105–4116 (2023).

27. Farahani, A. et al. Network spreading and local biological vulnerability in amyotrophic lateral sclerosis. Commun. Biol. 8, (2025).

28. Wainger, B. J. et al. Intrinsic membrane hyperexcitability of amyotrophic lateral sclerosis patient-derived motor neurons. Cell Rep. 7, 1–11 (2014).

29. Kshatri, A. S., Gonzalez-Hernandez, A. & Giraldez, T. Physiological Roles and Therapeutic Potential of Ca2+ Activated Potassium Channels in the Nervous System. Frontiers in Molecular Neuroscience vol. 11 Preprint at 10.3389/fnmol.2018.00258 (2018).

30. Pasniceanu, I. S. et al. Striatal neuron dysfunction in C9ORF72-FTD/ALS is driven by AIS and potassium channel dysregulation. Cell Rep. 45, (2026).

31. Joseph, B. J. et al. TDP-43-dependent mis-splicing of KCNQ2 triggers intrinsic neuronal hyperexcitability in ALS/FTD. Nat. Neurosci. 28, 2476–2492 (2025).

32. Stringer, R. N. & Weiss, N. Pathophysiology of ion channels in amyotrophic lateral sclerosis. Molecular Brain vol. 16 Preprint at 10.1186/s13041-023-01070-6 (2023).

33. Ma, X. R. et al. TDP-43 represses cryptic exon inclusion in the FTD–ALS gene UNC13A. Nature 2022 603:7899 603, 124–130 (2022).

34. Brown, A. L. et al. TDP-43 loss and ALS-risk SNPs drive mis-splicing and depletion of UNC13A. Nature 603, 131–137 (2022).

35. Dolphin, A. C. & Lee, A. Presynaptic calcium channels: specialized control of synaptic neurotransmitter release. Nature Reviews Neuroscience vol. 21 213–229 Preprint at 10.1038/s41583-020-0278-2 (2020).

36. Sirozh, O. et al. Nucleolar stress caused by arginine-rich peptides triggers a ribosomopathy and accelerates aging in mice. Mol. Cell 84, 1527–1540.e7 (2024).

37. Kasper, E. et al. Alzheimer’s Disease Co-Pathology and Cognitive Impairment in Amyotrophic Lateral Sclerosis. Ann. Neurol. 10.1002/ana.78227 (2026) doi:10.1002/ana.78227.

38. Wei, Z., Iyer, M. R., Zhao, B., Deng, J. & Mitchell, C. S. Artificial Intelligence-Assisted Comparative Analysis of the Overlapping Molecular Pathophysiology of Alzheimer’s Disease, Amyotrophic Lateral Sclerosis, and Frontotemporal Dementia. Int. J. Mol. Sci. 25, (2024).

39. Luan, W. et al. Synaptic changes contribute to persistent extra-motor behaviour deficits in amyotrophic lateral sclerosis. Acta Neuropathologica Communications 14, (2026).

40. Niven, E. et al. Validation of the Edinburgh Cognitive and Behavioural Amyotrophic Lateral Sclerosis Screen (ECAS): A cognitive tool for motor disorders. Amyotroph. Lateral Scler. Frontotemporal Degener. 16, 172–179 (2015).

41. Gillingwater, T. H. et al. Delayed synaptic degeneration in the CNS of Wlds mice after cortical lesion. Brain 129, 1546–1556 (2006).

42. Kay, K. R. et al. Studying synapses in human brain with array tomography and electron microscopy. Nat. Protoc. 8, 1366–1380 (2013).

